# Cardiac Cell Type-Specific Gene Regulatory Programs and Disease Risk Association

**DOI:** 10.1101/2020.09.11.291724

**Authors:** James D. Hocker, Olivier B. Poirion, Fugui Zhu, Justin Buchanan, Kai Zhang, Joshua Chiou, Tsui-Min Wang, Xiaomeng Hou, Yang E. Li, Yanxiao Zhang, Elie N. Farah, Allen Wang, Andrew D. McCulloch, Kyle J. Gaulton, Bing Ren, Neil C. Chi, Sebastian Preissl

## Abstract

**Background:** *Cis*-regulatory elements such as enhancers and promoters are crucial for directing gene expression in the human heart. Dysregulation of these elements can result in many cardiovascular diseases that are major leading causes of morbidity and mortality worldwide. In addition, genetic variants associated with cardiovascular disease risk are enriched within *cis*-regulatory elements. However, the location and activity of these *cis*-regulatory elements in individual cardiac cell types remains to be fully defined.

**Methods:** We performed single nucleus ATAC-seq and single nucleus RNA-seq to define a comprehensive catalogue of candidate *cis*-regulatory elements (cCREs) and gene expression patterns for the distinct cell types comprising each chamber of four non-failing human hearts. We used this catalogue to computationally deconvolute dynamic enhancers in failing hearts and to assign cardiovascular disease risk variants to cCREs in individual cardiac cell types. Finally, we applied reporter assays, genome editing and electrophysiogical measurements in *in vitro* differentiated human cardiomyocytes to validate the molecular mechanisms of cardiovascular disease risk variants.

**Results:** We defined >287,000 candidate *cis*-regulatory elements (cCREs) in human hearts at single-cell resolution, which notably revealed gene regulatory programs controlling specific cell types in a cardiac region/structure-dependent manner and during heart failure. We further report enrichment of cardiovascular disease risk variants in cCREs of distinct cardiac cell types, including a strong enrichment of atrial fibrillation variants in cardiomyocyte cCREs, and reveal 38 candidate causal atrial fibrillation variants localized to cardiomyocyte cCREs. Two such risk variants residing within a cardiomyocyte-specific cCRE at the *KCNH2/HERG* locus resulted in reduced enhancer activity compared to the non-risk allele. Finally, we found that deletion of the cCRE containing these variants decreased *KCNH2* expression and prolonged action potential repolarization in an enhancer dosage-dependent manner.

**Conclusions:** This comprehensive atlas of human cardiac cCREs provides the foundation for not only illuminating cell type-specific gene regulatory programs controlling human hearts during health and disease, but also interpreting genetic risk loci for a wide spectrum of cardiovascular diseases.

## INTRODUCTION

Disruption of gene regulation is an important contributor to cardiovascular disease, the leading cause of morbidity and mortality worldwide^1^. *Cis*-regulatory elements such as enhancers and promoters are crucial for regulating gene expression^2–4^. Mutations in transcription factors and chromatin regulators can result in heart disease^5, 6^, and genetic variants associated with risk of cardiovascular disease are enriched within annotated candidate *cis*-regulatory elements (cCREs) in the human genome^7^. However, a major barrier to understanding the genetic and molecular basis of cardiovascular diseases is the paucity of maps and tools to interrogate gene regulatory programs in the distinct cell types of the human heart. Recent single cell/nucleus RNA-seq^8–10^ and spatial transcriptomic^11^ studies have revealed gene expression patterns in distinct cardiac cell types across developmental and adulthood stages in the human heart, including some which display gene expression patterns that are cardiac chamber/region-specific^9, 10^. However, the transcriptional regulatory programs responsible for cell type-specific and chamber-specific gene expression, and their potential links to non-coding risk variants for cardiovascular diseases and traits, remain to be fully defined.

Candidate *cis*-regulatory elements (cCREs) have been annotated in the human genome with the use of ChIP-seq, DNase-Seq, ATAC-seq, GRO-seq, etc. in a broad spectrum of human tissues including in bulk heart tissues and in purified cardiomyocytes^2–4, 12–15^. These maps have provided important insights into dynamic gene regulation during heart failure^14–16^ and begun to shed light on the function of non-coding cardiovascular disease variants^7, 12, 15^. However, major limitations of these studies including their focus on particular chambers/regions of the heart and failure to interrogate *cis*-regulatory elements across all distinct cardiac cell types, have restricted their utility in understanding how specific gene regulatory mechanisms may impact distinct cell types and regions of human hearts in health and disease. Although recent single cell genomic tools provide the opportunity to interrogate *cis*-regulatory elements at single cell resolution^16–20^, their application to mammalian hearts has been limited to a few adult and fetal mouse hearts^20, 21^. Thus, to comprehensively investigate *cis*-regulatory elements in the specific cell types of the human heart, we profiled chromatin accessibility in ∼80,000 heart cells using single nucleus ATAC-seq (snATAC-seq)^17, 18^ and created a comprehensive cardiac cell atlas of cCREs annotated by cell type and putative target genes. Integration of these data with single nucleus RNA-seq datasets from matched specimens revealed gene regulatory programs in nine major cardiac cell types. Using this human cardiac cCRE atlas, we further observed the remodeling of cell type-specific candidate enhancers during heart failure and the enrichment of cardiovascular disease-associated genetic variants in cCREs of specific cell types. Finally, we showed that a cardiomyocyte-specific enhancer harboring risk variants for atrial fibrillation is necessary for cardiomyocyte *KCNH2* expression and regulation of cardiac action potential repolarization.

## RESULTS

### Single nucleus analysis of chromatin accessibility and transcriptome in adult human hearts

To assess the accessible chromatin landscape of distinct cardiovascular cell types, we performed snATAC-seq^17^, also known as sciATAC-seq^18^, on all cardiac chambers from four adult human hearts without known cardiovascular disease (Supplemental Table I). We obtained accessible chromatin profiles for 79,515 nuclei, with a median of 2,682 fragments mapped per nucleus (Figure 1A, B, Supplemental Figure I, Supplemental Table II). We also performed single nucleus RNA-seq (snRNA-seq) for a subset of the above heart samples to complement the accessible chromatin data and obtained 35,936 nuclear transcriptomes, with a median of 2,184 unique molecular identifiers (UMIs) and 1,286 genes detected per nucleus (Figure 1A, C, Supplemental Figure II-A-F, Supplemental Table III). Using SnapATAC^22^ and Seurat^23^, we identified nine clusters from snATAC-seq (Figure 1B) and twelve major clusters from snRNA-seq (Figure 1C, Supplemental Figure II-G, H), which were annotated based on chromatin accessibility at promoter regions or expression of known lineage-specific marker genes, respectively^9, 10^ (Figure 1D, E, Supplemental Table IV). For example, chromatin accessibility and gene expression of atrial and ventricular cardiomyocyte markers such as *NPPA* and *MYH7*^24^ were used to classify these two cardiomyocyte subtypes (Figure 1D, E). Although gene expression patterns of lineage markers strongly correlated with accessibility at promoter regions across annotated cell types (Figure 1F) and single cell integration analysis^23^ revealed 93% concordance in annotation between snATAC-seq and snRNA-seq datasets (Supplemental Figure III, Supplemental Table III), some cellular sub-types identified from snRNA-seq including endocardial cells and myofibroblasts were not detected by snATAC-seq (Figure 1F). Additionally, atrial and ventricular cardiomyocyte nuclei from the left and right regions of the heart could be further clustered by transcriptome but not chromatin accessibility (Supplemental Figure II-I, J). We noted that cell type composition varied significantly between biospecimens and donors, highlighting the importance of single cell approaches to limit biases due to cell proportion differences in bulk assays (Supplemental Figure IV, Supplemental Tables II and III). In summary, we identified and annotated cardiac cell types using both chromatin accessibility and nuclear transcriptome profiles.

**Figure 1:**
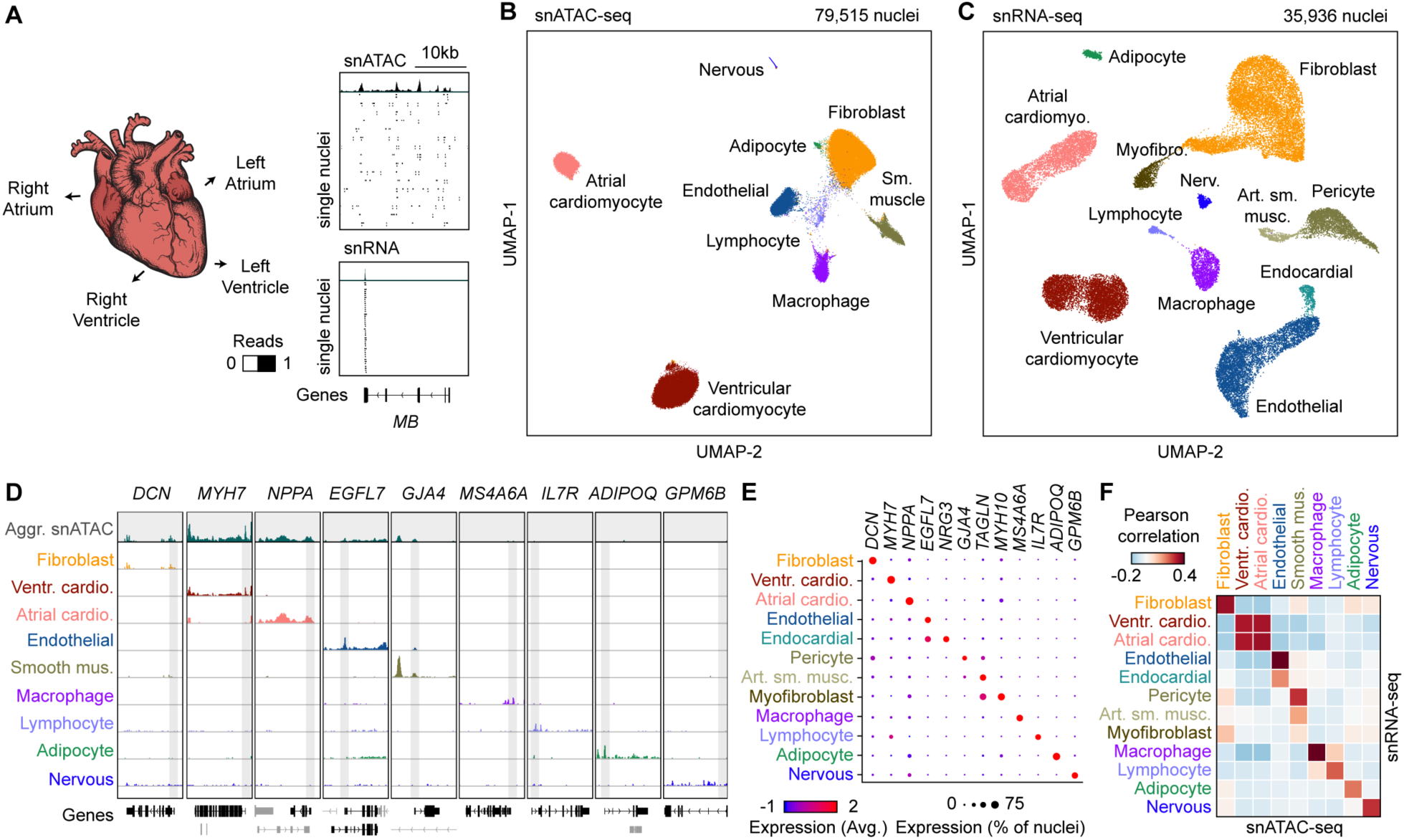
Single nucleus chromatin accessibility and transcriptome profiling of human hearts. **A)** snATAC-seq and snRNA-seq were performed on nuclei isolated from cardiac chambers from four human donors without cardiovascular pathology. snATAC-seq: n = 4 (left ventricle), n = 4 (right ventricle), n = 3 (left atrium), n = 2 (right atrium), snRNA-seq: n = 2 (left ventricle), n = 2 (right ventricle), n = 2 (left atrium), n = 1 (right atrium). **B)** Uniform manifold approximation and projection (UMAP)^59^ and clustering analysis of snATAC-seq data reveals nine clusters. Each dot represents a nucleus colored by cluster identity. **C)** Uniform manifold approximation and projection (UMAP)^59^ and clustering analysis of snRNA-seq data reveals 12 major clusters. Each dot represents a nucleus colored by cluster identity. Nerv. = Nervous. Art. sm. musc. = arterial smooth muscle. **D)** Genome browser tracks^60^ of aggregate chromatin accessibility profiles at selected representative marker gene examples for individual clusters and for all nuclei pooled together into an aggregated heart dataset (top track, grey). Black genes below tracks represent the indicated marker genes, non-marker genes are greyed. **E)** Dot plot illustrating expression of representative marker gene examples in individual snRNA-seq clusters. **F)** Heatmap illustrating the correlation between clusters defined by chromatin accessibility and transcriptomes. Pearson correlation coefficients were calculated between chromatin accessibility at cCREs within 2 kbp of annotated promoter regions^61^ and expression of the corresponding genes for each cluster.

### Identification of candidate *cis*-regulatory elements (cCREs) in distinct cell types of the human heart

To discover the cCREs in each cell type of the human heart, we aggregated snATAC-seq data from nuclei comprising each cell cluster individually and determined accessible chromatin regions with MACS2^25^. We then merged the peaks from all nine cell clusters into a union of 287,415 cCREs, which covered 4.7% of the human genome (Figure 2A, Supplemental Table V). 67.0% of the cCREs identified in the current study overlapped previously annotated cCREs from a broad spectrum of human tissues and cell lines^26, 27^ (Supplemental Figure V-A), and the union of heart cCREs captured 98.6% and 95.4% of candidate human heart enhancers reported in two previous bulk studies^12, 14^ (Supplemental Figure V-B, C). Furthermore, 75% of cCREs in the union were at least 2 kbp away from annotated promoter regions, and 19,447 displayed high levels of cell type-specificity (Figure 2B, Supplemental Table VI). Gene ontology analysis^28^ revealed that these cell type-specific cCREs were proximal to genes involved in relevant biological processes, including collagen fibril organization for cardiac fibroblast-specific cCREs (K1), and myofibril organization for ventricular cardiomyocyte-specific cCREs (K2, Figure 2C, Supplemental Table VII). Employing chromVAR^29^ (Supplemental Table VIII) and HOMER^30^ (Supplemental Table IX), we detected cell type-dependent enrichment for 231 transcription factor binding signatures, such as MEF2A/B, NKX2.5, and THR-β sequence motifs in cardiomyocyte-specific cCREs and TCF21 motifs in cardiac fibroblast-specific cCREs (Figure 2D, E). To discover the transcription factors that may bind to these sites, we combined corresponding snRNA-seq data with sequence motif enrichments to correlate expression of these transcription factors with motif enrichment patterns across cell types (Figure 2F). As an example, we found strong enrichment of the binding motif for the macrophage transcription factor SPI1/PU.1^31^ in macrophage-specific cCREs, and *SPI1* was exclusively expressed in macrophages (Figure 2F, Supplemental Tables IV and X). In addition, we observed that transcription factor family members were expressed in cell type-specific combinations. For instance, while GATA family members displayed similar motif enrichment patterns across sets of cell type-specific cCREs, we discovered that endothelial cells and cardiac fibroblasts expressed *GATA2* and *GATA6*, respectively, whereas cardiomyocytes expressed both *GATA4* and *GATA6,* and endocardial cells expressed *GATA2*, *GATA4*, and *GATA6* (Figure 2F, Supplemental Tables IV and X). In summary, these results establish a resource of candidate *cis*-regulatory elements for interrogation of cardiac cell type-specific gene regulatory programs.

**Figure 2:**
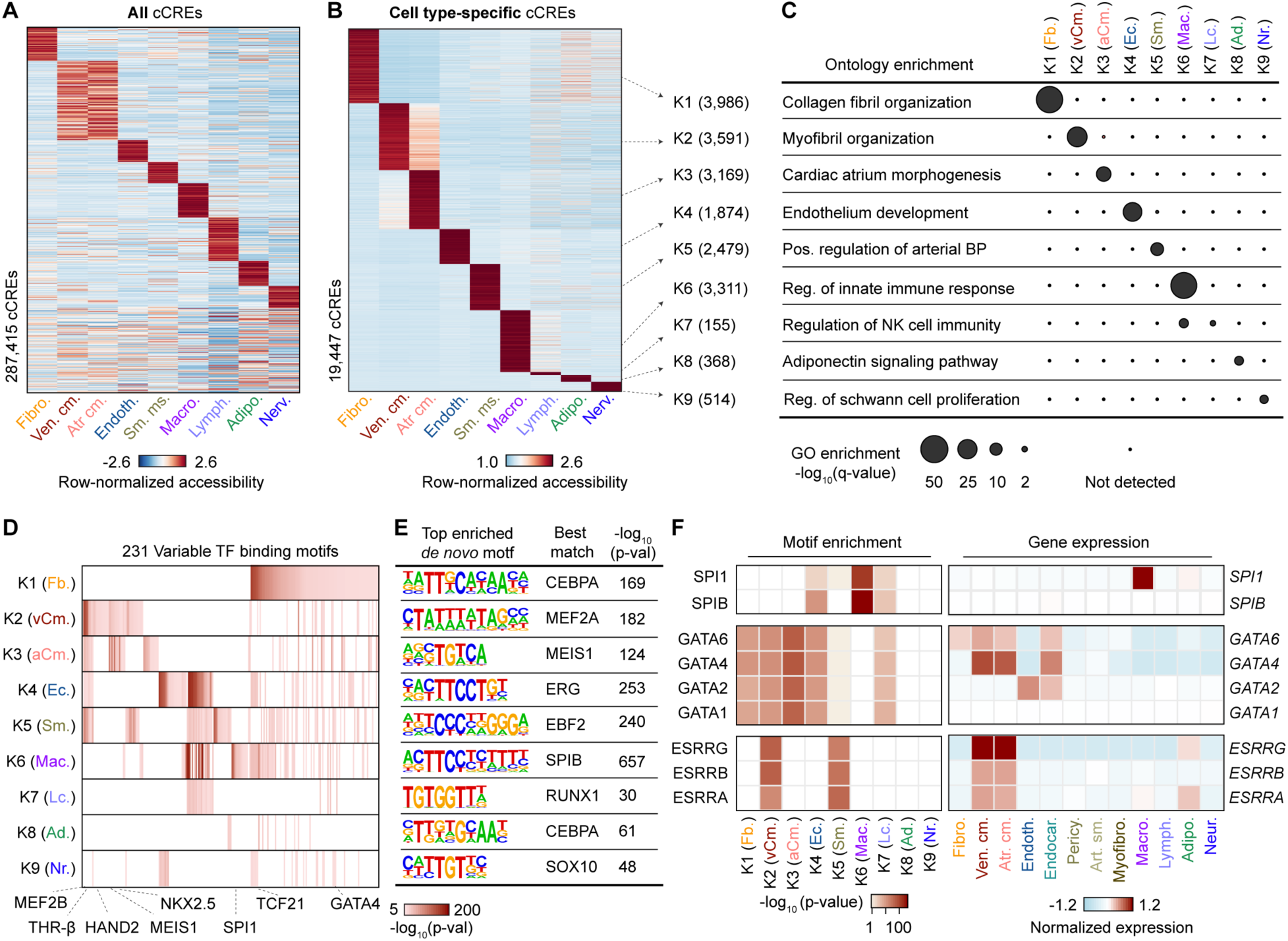
Characterization of gene regulatory programs in cardiac cell types. **A)** Heatmap illustrating row-normalized chromatin accessibility values for the union of 287,415 cCREs. K-means clustering was performed to group cCREs based on relative accessibility patterns. **B)** Heatmap showing row-normalized chromatin accessibility of 19,447 cell type-specific cCREs (FDR < 0.01 after Benjamini-Hochberg correction; log_2_(fold change) > 0). K-means clustering was performed to group cCREs based on relative accessibility patterns. Number of cCREs per K can be found in brackets. **C)** GREAT ontology analysis^28^ of cell type-specific cCREs. Q-value for enrichment indicates Bonferroni adjusted p-value. **D, E)** Transcription factor motif enrichment^30^ for known (**D**) and *de novo* motifs (**E**) within cell type-specific cCREs. The heatmap in (**D**) shows motifs with enrichment p-value <10^-5^ in at least one cluster. For *de novo* transcription factor motifs (**E**) the best matches for the top motifs are displayed. Statistical test for motif enrichment: hypergeometric test. P-values were not corrected for multiple testing. **F)** Combination of transcription factor motif enrichment and gene expression shows cell type-specific roles for members of transcription factor families. Displayed are heatmaps for known motif enrichment in cell type-specific cCREs (left) and gene expression across clusters (right). (Fb. = Fibroblast, vCm. = Ventricular Cardiomyocyte, aCm. = Atrial Cardiomyocyte, Ec. = Endothelial, Sm. = Smooth Muscle, Mac. = Macrophage, Lc. = Lymphocyte, Ad. = Adipocyte, Nr. = Nervous).

### Cardiac cell type-specific gene regulatory programs implicated in chamber-specific structure and function

Each cardiac chamber performs a unique role that is crucial to system-level heart function^32^. To investigate the gene regulatory programs underlying chamber-specific gene expression and cellular functions in distinct cardiac cell types, we tested cCREs for differential accessibility across five of the most abundant cell types of the heart: cardiomyocytes, cardiac fibroblasts, endothelial cells, smooth muscle cells, and macrophages. We discovered 16,451 differentially accessible (DA) cCREs between pooled atria and ventricles, the majority of which were detected in cardiomyocytes (Figure 3A-C, Supplemental Table X). Specifically, 11,159 cCREs displayed differential accessibility between right atrium and right ventricle and 12,962 cCREs exhibited differential accessibility between left atrium and left ventricle (Supplemental Figure VI-A-C, Supplemental Table X). Comparing the left and right sides of the heart, we identified 101 DA cCREs between the right and left ventricle (Supplemental Figure VI-D), and 2,687 DA cCREs between left and right atria, which in contrast to comparisons between atria and ventricles were found primarily in cardiac fibroblasts (Supplemental Figure VI-E, Supplemental Table X).

**Figure 3:**
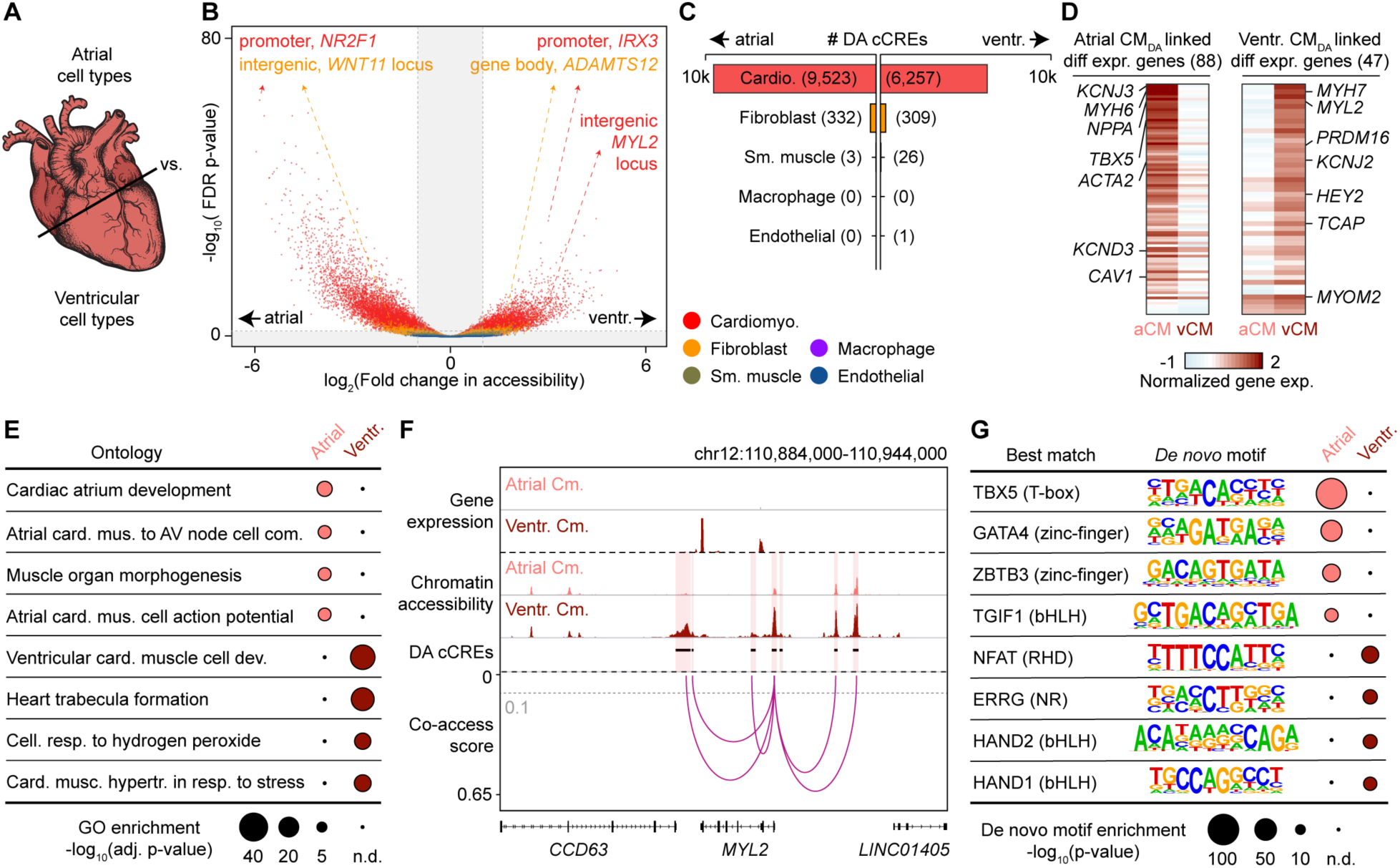
Cardiomyocyte cCREs display chamber-dependent differences in chromatin accessibility. **A)** Scheme for comparison of major cell types across heart chambers. All atrial as well as all ventricular datasets were combined, and corresponding cell types compared. **B)** Volcano plot showing differentially accessible (DA) candidate *cis*- regulatory elements (cCREs) in each cell type between atria and ventricles. Each dot represents a cCRE and the color indicates the cell type. cCREs with log_2_(fold change) > 1 and FDR < 0.05 after Benjamini-Hochberg correction (outside the shaded area) were considered as DA. **C)** DA cCREs between atria and ventricles were detected almost exclusively in cardiomyocytes and fibroblasts. The numbers of DA cCREs are listed in brackets. **D)** Heatmaps showing normalized gene expression levels of differentially expressed genes between atrial (aCM) and ventricular cardiomyocytes (vCM) that were linked by co-accessibility to distal DA cCREs that were more accessible in atrial cardiomyocytes (Atrial CM_DA_) or ventricular cardiomyocytes (Ventr. CM_DA_), respectively. **E)** GREAT ontology analysis^28^ of DA cCREs between atrial and ventricular cardiomyocytes. P-values shown are Bonferroni adjusted (n.d.: not detected). **F)** Genome browser tracks^60^ showing chromatin accessibility and gene expression in atrial and ventricular cardiomyocytes as well as DA cCREs that were co-accessible with the promoter of *MYL2*. Grey dotted line indicates co-accessibility threshold (> 0.1). Co-accessible DA cCREs are indicated by a red shaded box and the promoter region of *MYL2* is indicated by a grey shaded box. **G)** Transcription factor motif enrichment analysis^30^ of DA cCREs between atrial and ventricular cardiomyocytes. The best matches for the top *de novo* motifs (score > 0.7) are shown. Statistical test for motif enrichment: hypergeometric test. P-values were not corrected for multiple testing (n.d.: not detected).

Utilizing co-accessibility analysis^33^ to link distal DA cCREs (∼88% of all DA cCREs) to their putative target genes (Supplemental Table XI, median distance: 88.7 kbp), we observed that distal DA cCREs in cardiomyocytes between atria and ventricles were associated with chamber-specific gene expression of their putative target genes (Figure 3D, Supplemental Figure VI-B-E, Supplemental Table XII), and genes near these DA cCREs were enriched for chamber-specific biological processes (Figure 3E, Supplemental Figure VI-B-E, Supplemental Table XIII). Specifically, distal DA cCREs with higher accessibility in atrial cardiomyocytes were associated with genes such as *PITX2,* a transcriptional regulator of cardiac atrial development, as well as the ion channel subunit *SCN5A* which regulates cardiomyocyte action potential (Figure 3E, Supplemental Table XIII). Furthermore, we found distal DA cCREs with higher accessibility in atrial cardiomyocytes at the *HAMP* gene locus, which encodes a key regulator of ion homeostasis and was recently described as a potential novel cardiac gene in the right atrium by single nucleus transcriptomic analysis^9, 10^. Conversely, genes near distal DA cCREs with higher accessibility in ventricular cardiomyocytes were enriched for biological processes such as trabecula formation and ventricular cardiac muscle cell differentiation. For example, several distal DA cCREs with increased accessibility in ventricular cardiomyocytes compared to atrial cardiomyocytes were linked to the promoter region of *MYL2*, which encodes the ventricular isoform of myosin light chain 2^34^ (Figure 3F, Supplemental Table IV), a regulator of ventricular cardiomyocyte sarcomere function.

Additionally, analysis of distal DA cCREs in cardiac fibroblasts revealed that putative target genes were involved in distinct biological processes between right and left atria. In particular, we found that DA cCREs with higher accessibility in right atrial cardiac fibroblasts were proximal to genes involved in heart development, heart growth, and tube development, whereas DA cCREs with higher accessibility in left atrial cardiac fibroblasts were adjacent to genes involved in biological processes such as wound healing and vasculature development (Supplemental Figure VI-E, Supplemental Table XIII). We further found a cardiac fibroblast-specific DA cCRE with higher accessibility in left atria at the fibrinogen *FN1* gene locus, potentially indicating a more activated fibroblast state^9, 35^. Supporting these findings, we identified several other DA cCREs with higher accessibility in left atrial cardiac fibroblasts adjacent to genes involved in generation of extracellular matrix (ECM) such as *MMP2* and *FBLN2* (Supplemental Table XIII). These observations are consistent with previous findings that a higher fraction of ECM is produced in fibroblasts of the left atrium^9^.

Using motif enrichment analysis, we inferred candidate transcriptional regulators involved in chamber-specific cellular specialization, including TBX5, GATA4, and TGIF1 for atrial cardiomyocytes, and NFAT, ERRG, HAND1, and HAND2 for ventricular cardiomyocytes (Figure 3G, Supplemental Table XIV). While the TBX5 DNA binding motif was strongly enriched in both right and left atrial cardiomyocyte DA cCREs, the NFAT5 motif ranked highest in left ventricular cardiomyocyte DA cCREs and the TBX20 motif was strongly enriched in right ventricular cardiomyocyte DA cCREs (Supplemental Figure VI-B, C, Supplemental Table XIV). Furthermore, cardiac fibroblast DA cCREs with higher accessibility in the right atrium were enriched for the binding motif of forkhead transcription factors (Supplemental Figure VI-E), whereas cardiac fibroblast DA cCREs with higher accessibility in the left atrium were enriched for the homeobox transcription factor CUX1 motif (Supplemental Figure VI-E, Supplemental Table XIV). Altogether, we identified cCREs and candidate transcription factors associated with specific cardiac chambers, particularly within cardiomyocytes and cardiac fibroblasts.

### Cell type specificity of candidate enhancers associated with heart failure

Recent large-scale studies profiling the H3K27ac histone modification in human hearts have uncovered candidate enhancers associated with heart failure^14, 16^. However, because these studies either examined heterogeneous bulk heart tissue^14, 16^ or focused solely on enriched cardiomyocytes^15^, it remains unclear what role, if any, additional cardiac cell types and cCREs may contribute to heart failure pathogenesis. Using our cell atlas of cardiac cCREs, we revealed the cell type specificity of candidate enhancers showing differential H3K27ac signal strength between human hearts from healthy donors and donors with dilated cardiomyopathy (heart failure)^14^ (Figure 4, Supplemental Figure VII). We observed that a large fraction of candidate enhancers that displayed increased activity (45%) during heart failure were accessible primarily in cardiac fibroblasts (Figure 4A, K2-4_up,_ Supplemental Table XV), whereas a majority of those exhibiting decreased activity (67%) were accessible primarily in cardiomyocytes (Figure 4B, K1-3_down_, Supplemental Table XV). Candidate enhancers with increased activity in cardiac fibroblasts were proximal to genes involved in extracellular matrix organization and connective tissue development (Figure 4A, K2-4_up_, Supplemental Table XVI), whereas those exhibiting decreased activity in cardiomyocytes were proximal to genes involved in regulation of heart contraction and cation transport (Figure 4B, K1-3_down_, Supplemental Table XVI). For example, several of these cardiac fibroblast candidate enhancers were present at loci encoding the extracellular matrix proteins lumican (*LUM*) and decorin (*DCN*) and co-accessible with the promoters of these genes (Figure 4C). Consistent with these findings, both genes were primarily expressed in cardiac fibroblasts (Supplemental Table IV), and *LUM* has been reported to exhibit increased expression in failing hearts compared to control hearts^14^. On the other hand, several cardiomyocyte candidate enhancers displaying decreased activity in heart failure were co-accessible with the promoter region of *IRX4* (Figure 4D), which encodes a ventricle-specific transcription factor^36^ and is specifically expressed in cardiomyocytes of the left ventricle (Supplemental Table IV).

**Figure 4:**
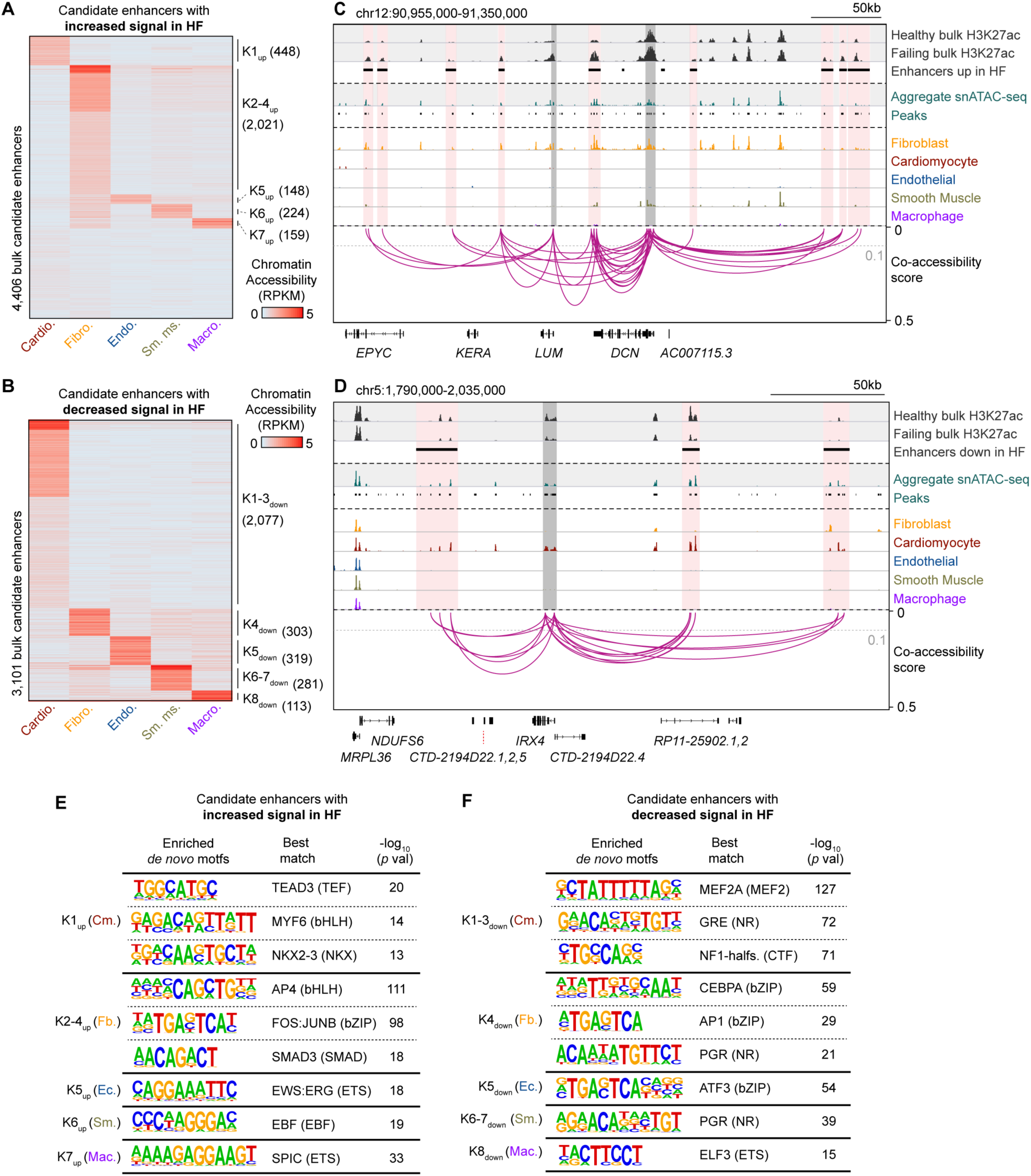
Cell type specificity of candidate enhancers associated with heart failure. **A)** Cell type-specificity of 4,406 candidate enhancers with increased H3K27ac signal in failing left ventricles^14^. Heatmap displays cell type-resolved chromatin accessibility RPKM (reads per kilobase per million mapped reads) values for cell types from left ventricular snATAC-seq datasets. Candidate enhancers were grouped based on chromatin accessibility patterns across cell clusters using K-means. **B)** Cell type-specificity of 3,101 candidate enhancers with decreased H3K27ac signal in failing left ventricles^14^. **C)** Genome browser tracks^60^ showing several candidate enhancers with increased activity during heart failure (HF) that were primarily accessible in fibroblasts and co-accessible with the promoters of *LUM* and/or *DCN.* For visualization, linkages between cCREs within candidate enhancers and all gene promoters are shown (co-accessibility > 0.1, grey dotted line). Candidate enhancers co-accessible with gene promoters are indicated by red shaded boxes and promoter regions are indicated by grey shaded boxes. **D)** Genome browser tracks^60^ showing several bulk candidate enhancers with decreased activity in heart failure that were primarily accessible in cardiomyocytes and co-accessible with the promoter of *IRX4.* **E, F)** Transcription factor motif enrichment^30^ in the candidate enhancers with (**E**) increased and (**F**) decreased activity in failing left ventricles. Analysis was performed on the indicated K cluster(s) from panels (**A**) and (**B**) respectively. The best matches for selected *de novo* motifs (score > 0.7) are shown. Statistical test for motif enrichment: hypergeometric test. P-values were not corrected for multiple testing.

To identify potential transcription factors regulating these pathologic responses during heart failure, we performed motif enrichment analysis in cell type-specific subsets of disease-associated candidate enhancers (Supplemental Table XVII). For candidate enhancers exhibiting increased activity in heart failure, we identified enrichment of not only bHLH motifs such as AP4 in cardiac fibroblast candidate enhancers which matched previous bulk analysis^14^ (Figure 4E, K2-4_up_), but also TEAD3 and MYF6 motifs in cardiomyocyte candidate enhancers (Figure 4E, K1_up_). Conversely, for candidate enhancers displaying decreased activity in heart failure, we observed enrichment of nuclear receptor motifs such as glucocorticoid response element (GRE) in cardiomyocyte candidate enhancers, which is consistent with previous findings^14^ (Figure 4F, K1-3_down_), as well as other motifs which were not detected in bulk analyses, such as the bZIP transcription factor CEBPA for cardiac fibroblast candidate enhancers (Figure 4F, K4_down_). Thus, these results show that this cardiac cell atlas of cCREs may be used to assign disease-associated candidate enhancers from bulk assays to their affected cell types and infer transcriptional regulators involved in lineage-specific disease pathogenesis.

### Interpreting non-coding risk variants of cardiac diseases and traits

Non-coding genetic variants contributing to risk of complex diseases are enriched within cCREs in a cell type-specific fashion^20, 37–40^. To examine the enrichment of cardiovascular disease variants within cCREs active in specific cardiac cell types, we performed cell type-stratified LD (Linkage disequilibrium) score regression analysis^41^ using GWAS summary statistics for cardiovascular diseases^42–46^ (Figure 5A) and control traits (Supplemental Figure VIII, Supplemental Table XVIII). This analysis revealed significant enrichment of atrial fibrillation (AF)-associated variants in both atrial (Z = 3.25, FDR = 0.02) and ventricular cardiomyocyte cCREs (Z = 3.77, FDR = 0.01), varicose vein-associated variants in endothelial cell cCREs (Z = 3.44, FDR = 0.01), and nominal enrichment of coronary artery disease-associated variants in cardiac fibroblast cCREs (Z = 2.19, FDR = 0.20, Figure 5A).

**Figure 5:**
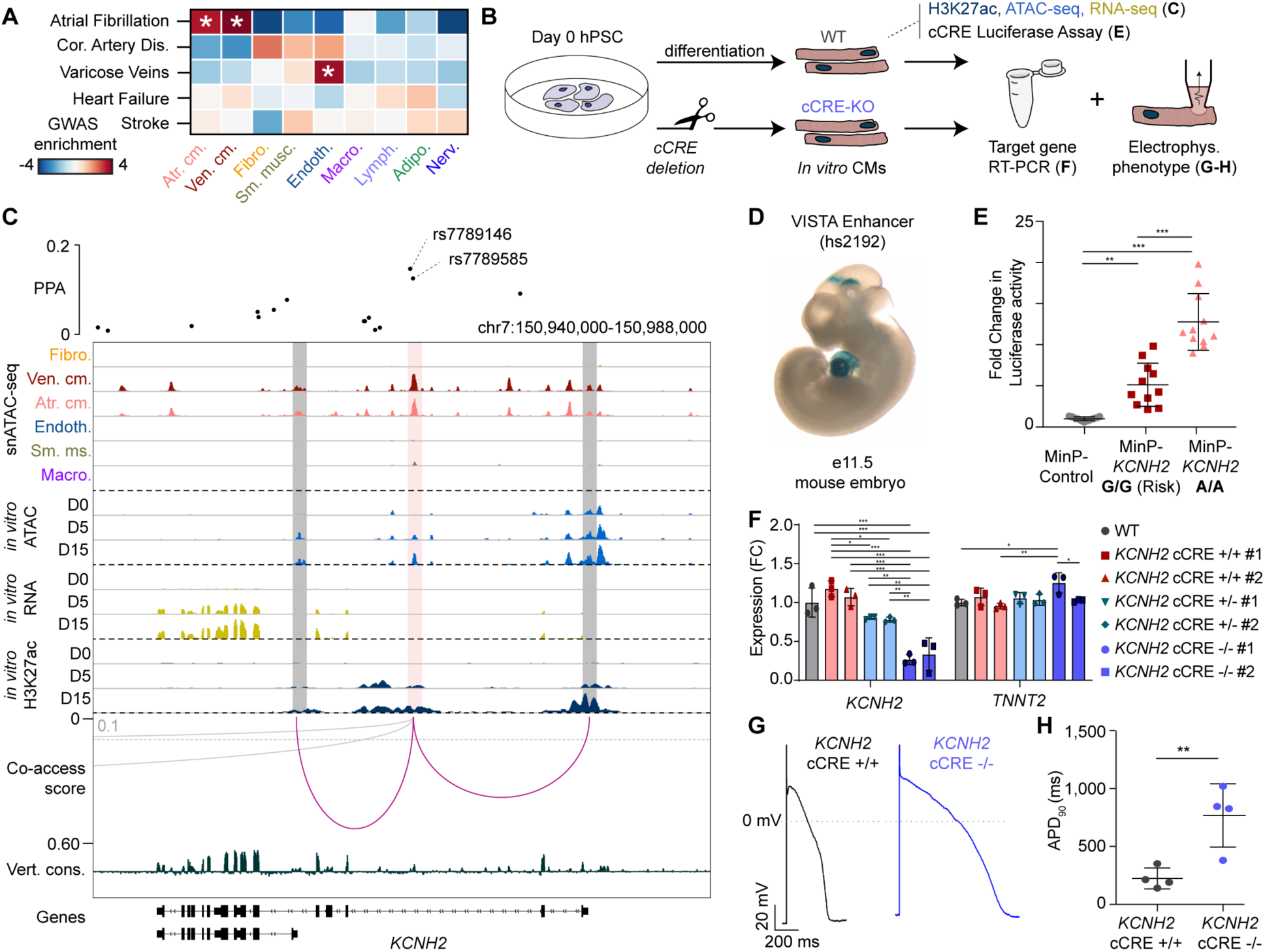
Identification and characterization of atrial fibrillation-associated variants at the *KCNH2* locus. **A)** Enrichment of risk variants associated with cardiovascular disease traits in genome wide association studies (GWAS) within cardiac cell type-resolved cCREs. Displayed are z-scores, and these scores were also used to compute one-sided p-values for enrichments that were corrected using the Benjamini Hochberg procedure for multiple tests. * = FDR < 0.05. **B)** Schematic of a cardiomyocyte differentiation model used to profile candidate enhancer dynamics, gene expression, and electrophysiologic phenotypes. hPSC = human pluripotent stem cell. **C)** Fine mapping^47^ and molecular characterization of two variants associated with atrial fibrillation (AF) in a cardiomyocyte cCRE co-accessible with promoter regions of *KCNH2*. Genome browser tracks^60^ display cell type-resolved chromatin accessibility and co-accessibility from snATAC-seq, as well as chromatin accessibility, H3K27ac signal, and gene expression during hPSC-cardiomyocyte differentiation timepoints. D0 = day 0, D5 = day 5, D15 = day15. For illustration, the co-accessibility track shows linkages between the AF variant-containing cCRE and annotated gene promoters (cutoff > 0.1, grey dotted line). The grey arc represents links to the promoter of *AOC1* which was not expressed. For the full locus see Supplemental Figure IX-A. PPA: Posterior probability of association^47^. **D)** Representative image of a transgenic mouse embryo showing LacZ reporter gene expression under control of a genomic region (hs2192, image downloaded from Vista database^50^, https://enhancer.lbl.gov/) that overlaps the variant-cCRE pair at the *KCNH2* locus. The picture for hs2192 was downloaded from the VISTA^50^ database. **E)** Dot plot illustrating results of a dual luciferase reporter assay for the AF variant-harboring cCRE at the *KCNH2* locus in D15 cardiomyocytes. The genotype for both rs7789146 and rs7789585 was either G (homozygous rs7789146-G / rs7789585-G; risk) or A (homozygous rs7789146-A / rs7789585-A; non-risk). Each dot represents one transfection (three independent experiments). Data are displayed as mean +/− SD. *** p < 0.001, ** p < 0.01 (one-way ANOVA and Tukey post hoc test). MinP: minimal promoter. **F)** Bar chart showing fold changes in *KCNH2* and *TNNT2* expression for D25 cardiomyocytes measured by qPCR after CRIPSR/Cas9-mediated deletion of the variant-cCRE pair at the *KCNH2* locus. Each dot represents one independent cardiomyocyte differentiation. Data are displayed as mean +/− SD. *** p < 0.001, ** p < 0.01, * p < 0.05, (one-way ANOVA and Tukey post hoc test); WT = unperturbed control, *KCNH2* cCRE +/+ #1 & #2 = no guide RNA control clones #1 and #2; *KCNH2* cCRE +/− #1 & #2 = Heterozygous enhancer deletion clones #1 & #2; *KCNH2* cCRE −/− #1 & #2 = Homozygous enhancer deletion clones #1 & #2. **G)** Exemplary traces of action potential recordings in hPSC-derived cardiomyocytes at D25-35 for a control clone (*KCNH2* cCRE +/+ #1, left) and a clone with enhancer deletion (*KCNH2* cCRE −/− #1, right). **H)** Dot blot showing the quantification of APD_90_ at 1 Hz pacing for 4 independent hPSC derived cardiomyocytes at D25-35 from a control clone (*KCNH2* cCRE +/+ #1) and an enhancer deletion clone (*KCNH2* cCRE −/− #1). ** p < 0.01 (unpaired two-sided t-test). APD_90_: action potential duration at 90% depolarization.

Next, to identify likely causal AF risk variants in cardiomyocyte cCREs, we first determined the probability that variants were causal for AF (Posterior probability of association, PPA) at 111 known loci using Bayesian fine-mapping^47^. We then intersected fine-mapped AF variants with cCREs and identified 38 variants with PPA > 10% in cardiomyocyte cCREs including previously reported variants at the *HCN4*^12^ and *SCN10A/SCN5A*^48^ loci (Supplemental Table XIX). We further prioritized AF variants for molecular characterization based on their overlap with cCREs that were primarily accessible in cardiomyocytes, evolutionarily conserved, co-accessible with promoters of genes expressed in cardiomyocytes and marked by H3K27ac in human pluripotent stem cell (hPSC)-derived cardiomyocytes^49^ during *in vitro* differentiation (Figure 5B). From this analysis, we discovered a cCRE in the second intron of the potassium channel gene *KCNH2* (*HERG*) which was co-accessible with the *KCNH2* promoter (Figure 5C) and harbored two variants, rs7789146 and rs7789585, with a combined PPA of 28% (Figure 5C, Supplemental Figure IX-A). This cCRE appeared to be activated during hPSC-cardiomyocyte differentiation as evidenced by an increase in H3K27ac signal that correlated with *KCNH2* expression (Figure 5C). Supporting its *in vivo* role in regulating gene expression in mammalian hearts, a genomic region (hs2192)^50^ containing this cCRE was previously shown to drive LacZ reporter expression in mouse embryonic hearts^50^ (Figure 5D).

### A cardiomyocyte enhancer of *KCNH2* is affected by non-coding risk variants associated with atrial fibrillation

To investigate whether these AF variants may affect enhancer activity and thereby regulate *KCNH2* expression and cardiomyocyte electrophysiologic function, we initially carried out reporter assays using a hPSC cardiomyocyte model system. Results from these studies confirmed that in D15 hPSC-cardiomyocytes, the *KCNH2* enhancer carrying the homozygous rs7789146-G/rs7789585-G AF risk allele displayed significantly weaker enhancer activity than when containing the non-risk variants (Figure 5E, Supplemental Figure IX-B), thus supporting the functional significance of these AF variants. We next used CRISPR/Cas9 genome editing strategies to remove the enhancer and performed qPCR and electrophysiologic assays to examine its role in *KCNH2* expression and function. Supporting the aforementioned findings, CRISPR/Cas9 genome deletion of this cCRE in hPSC-cardiomyocytes resulted in decreased *KCNH2* expression in an enhancer dosage-dependent manner (Figure 5F, Supplemental Figure IX-C). Similar to human cardiomyocytes with loss of *KCNH2* function due to mutations in the *KCNH2* coding sequence^51^ or gene knockdown^52^, cellular electrophysiologic studies demonstrated that these cCRE-deleted hPSC cardiomyocytes displayed a significantly prolonged action potential duration (Figure 5G, H), thus suggesting that cardiac repolarization abnormalities in atrial cardiomyocytes may lead to AF in an analogous manner to ventricular arrhythmias due to long QT syndrome^52^. Taken together, these results highlight the utility of this single cell atlas for assigning non-coding cardiovascular disease risk variants to distinct cell types and affected cCREs, and functionally interrogating how these variants may contribute to cardiovascular disease risk.

## DISCUSSION

The limited ability to interrogate cell type-specific gene regulatory programs in the human heart has been a major barrier for understanding molecular mechanisms of cardiovascular traits and diseases. Here, we report a cell type-resolved atlas of cCREs in the human heart, which was ascertained by profiling accessible chromatin in individual nuclei from all four chambers of multiple human hearts and includes both cell type-specific and heart chamber-specific cCREs. Furthermore, we characterized candidate *cis*-regulatory elements in different cardiac cell types in the human heart and delineated differences of gene regulatory programs underlying different regions/structures of the heart. In particular, we observed chamber-specific differences in chromatin accessibility between ventricles and atria as well as left and right atria but notably detected few differences between left and right ventricles. This finding is consistent with a recent single nucleus RNA-seq study in human hearts which found few differentially expressed genes between left and right ventricles^10^.

We further highlight the utility of this atlas of heart cCREs to provide new insight into aberrant gene regulation during cardiovascular pathology. To this end, we delineated the cell type-specificity of enhancers which were differentially active between healthy and failing heart tissue^14^ and identified additional transcription factors that may be involved in the pathogenesis of specific cell types during heart failure. Such cell type-specific analysis is particularly important in the context of heart failure because cellular composition can differ between diseased and control hearts^15, 53^. This change in cellular composition may in part explain the cell type bias that we observed between candidate enhancers exhibiting increased and decreased activity during heart failure (i.e. cardiac fibroblasts and cardiomyocytes, respectively). However, due to the large differences in H3K27ac signal, we suspect that measured changes in candidate enhancer activity could be due to a combination of both enhancer remodeling and shift in cell type composition. Thus, future studies profiling snATAC-seq and H3K27ac in parallel from the same cardiac sample or novel approaches to profile histone modifications in single nuclei^54, 55^ will provide greater insight into the extent of changes in chromatin accessibility and enhancer activity in individual cardiac cell types from diseased hearts.

Finally, we show how this atlas can be used to not only assign non-coding genetic variants associated with cardiovascular disease risk to cCREs in specific cardiac cell types, but also illuminate their cellular and molecular consequences. In particular, we discovered significant enrichment of AF-associated variants within cardiomyocyte cCREs and functionally interrogated one of these cCREs by demonstrating its role in regulating *KCNH2* expression and cardiomyocyte repolarization. Similar to electrophysiologic phenotypes of human cardiomyocytes exhibiting *KCNH2* loss of function^51, 52^, hPSC-cardiomyocytes harboring deletions of this cCRE displayed action potential prolongation, suggesting that cardiac repolarization abnormalities may contribute to atrial fibrillation, possibly through similar mechanisms as to how they may contribute ventricular arrhythmias^51^. On the other hand, we found only nominal enrichment of variants associated with coronary artery disease in fibroblasts and no enrichment of variants associated with heart failure in any cardiac cell type. These findings may reflect the heterogeneous etiologies of cardiovascular diseases and, in the case of heart failure, the limited number of currently known risk loci^42^. Future GWAS in large cohorts with detailed phenotyping, including biobanks such as the UK Biobank^56^ and the BioBank Japan Project^57^ and whole genome sequencing efforts such as the NHLBI Trans-Omics for Precision Medicine (TOPMed) program^58^, will help identify and refine disease association signals. Therefore, this atlas of cardiac cCREs will be a valuable resource for continued discovery of regulatory elements, target genes, and specific cell types that may be affected by non-coding cardiovascular genetic variants.

In summary, we created a human heart cell atlas of >287,000 cCREs, which may serve as a reference to further expand our knowledge of gene regulatory mechanisms underlying cardiovascular disease. To facilitate distribution of these data, we created a web portal at: http://catlas.org/humanheart. Integrating this resource with genomic and epigenomic clinical cardiac datasets, we built a systematic framework to interrogate how *cis*-regulatory elements and genetic variants might contribute to cardiovascular diseases such as heart failure or atrial fibrillation. Overall, such information will have great potential to provide new insight into the development of future cardiac therapies that are tailored to affected cell types and thus optimized for treating specific cardiovascular diseases.

## Supporting information

Supplemental_Table_III

Supplemental_Table_VII

Supplemental_Table_VIII

Supplemental_Table_XII

Supplemental_Table_XIII

Supplemental_Table_XIV

Supplemental_Table_XIX

Supplemental_Table_XVI

Supplemental_Table_XVII

Supplemental_Table_XVIII

Supplemental_Table_XXI

## ACKNOWLEDGEMENTS

We thank B. Li for bioinformatics support. We thank K. Jepsen and the UCSD IGM Genomics Center for sequencing the snRNA-seq libraries. We thank the QB3 Macrolab at UC Berkeley for purification of the Tn5 transposase.

## AUTHOR CONTRIBUTIONS

J.D.H., S.P., N.C.C., and B.R. conceived the project. J.D.H. and J.B. carried out snATAC-seq and snRNA-seq library preparation with help from X.H. F.Z. performed luciferase assay, CRISPR–Cas9 knockout, *in vitro* cardiomyocyte differentiation and quantitative PCR of the corresponding cell lines. T.W. and F.Z. performed action potential measurement. J.D.H., O.P., J.B., K.Z., J.C., Y.L., and S.P. performed data analysis. O.P. created the web portal. X.H., E.F., Y.Z., A.W., A.D.M., K.J.G., and N.C.C. contributed to experimental design and computational analyses. J.D.H., S.P., N.C.C., and B.R. wrote the manuscript. All authors edited and approved the manuscript.

## SUPPLEMENTAL FIGURES

**Supplemental Figure I:**
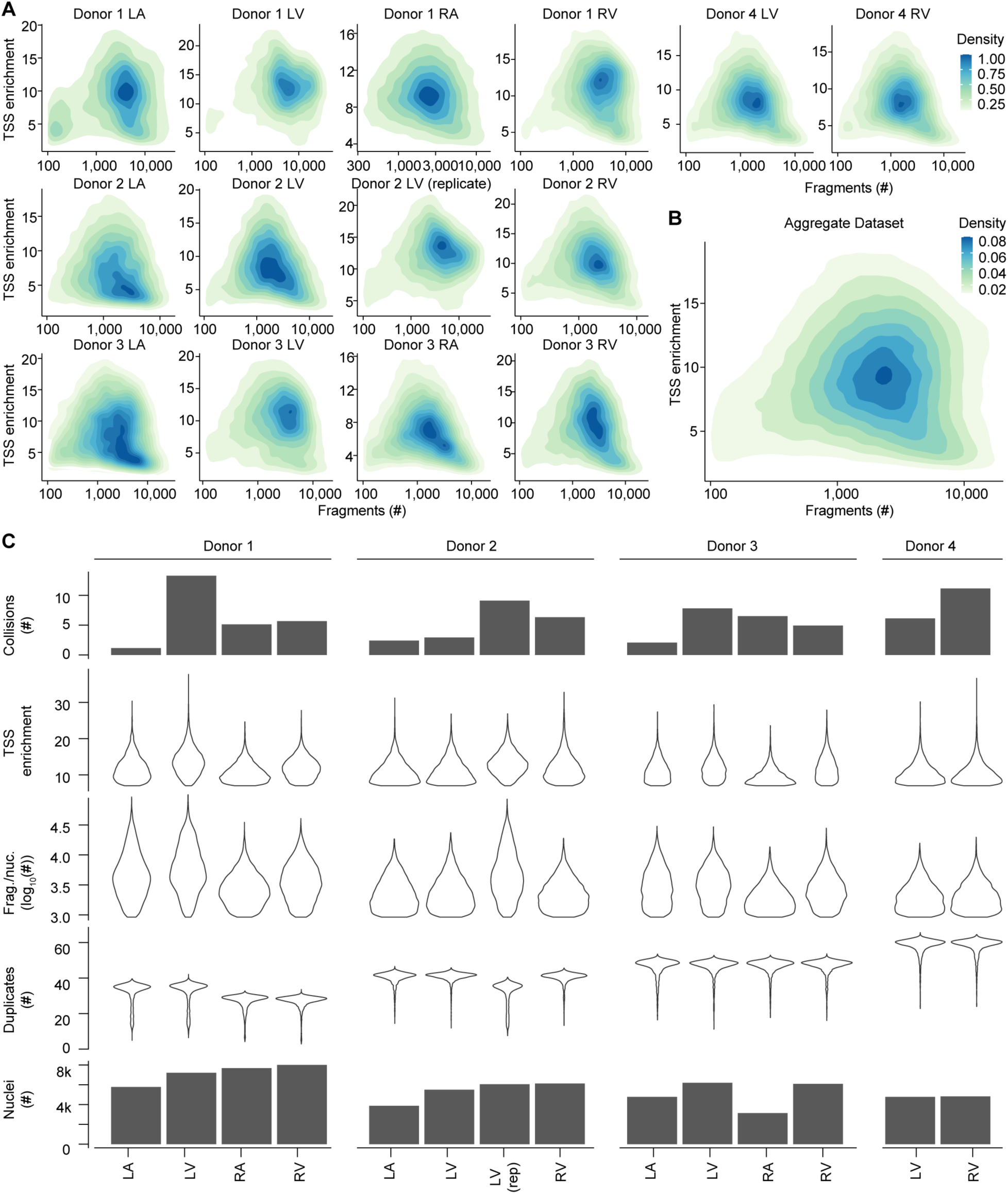
Quality control for snATAC-seq datasets. **A)** Density plots showing enrichment of fragments at transcription start sites (TSS enrichment) versus number of fragments per nucleus for each dataset. **B)** Density plot of TSS enrichment versus number of fragments for all datasets combined. **C)** Percentage of barcode collisions identified as heterotypic cell type collisions by Scrublet^62^ (top row), TSS enrichment (second row), fragments per nucleus (third row), duplicate read percentage (fourth row), and number of nuclei passing quality control (bottom row) for each snATAC-seq dataset.

**Supplemental Figure II:**
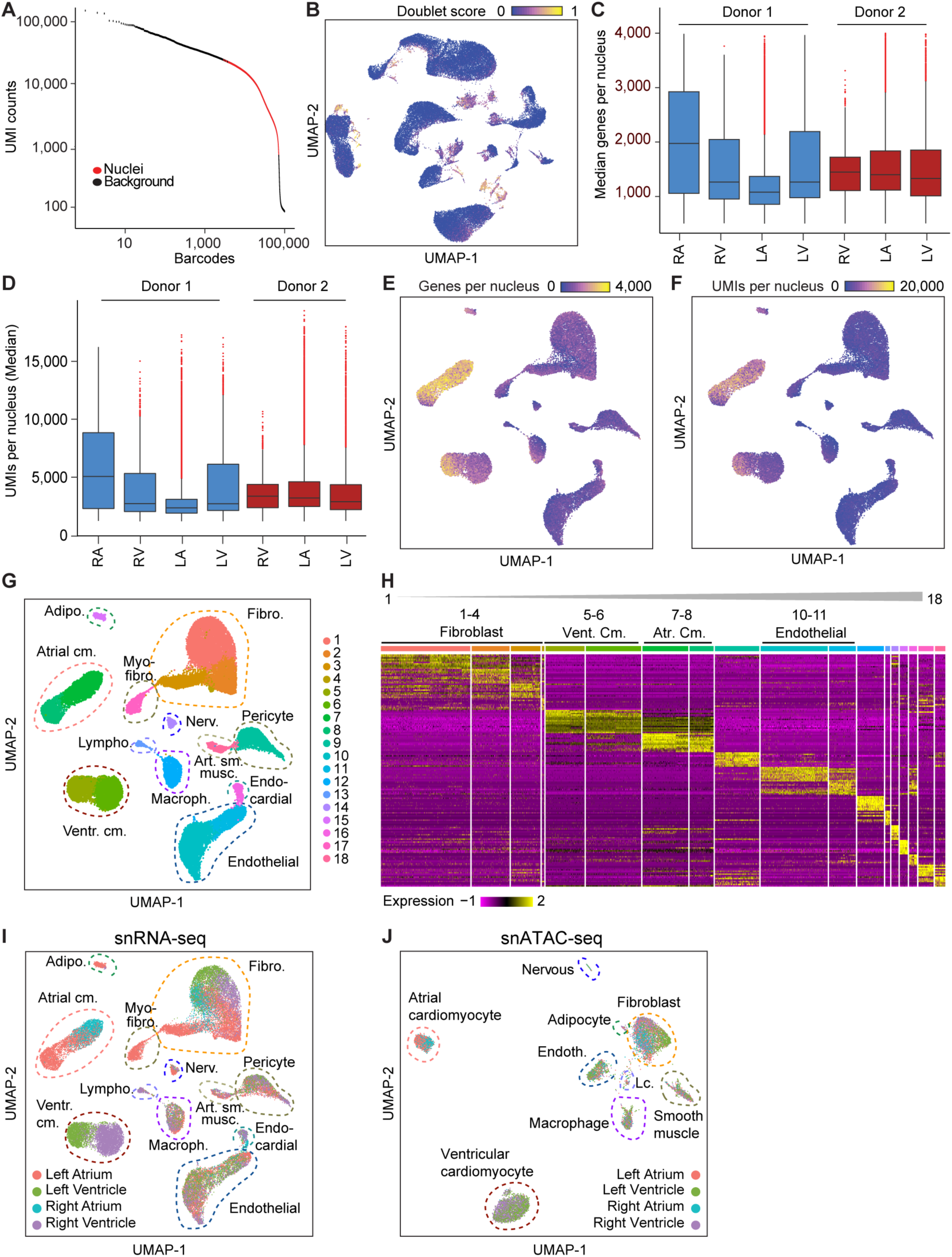
Quality control for snRNA-seq datasets and annotation of snRNA-seq clusters. **A)** Distribution of barcodes by unique molecular identifier (UMI) counts for nuclei (red; passing quality control) and background (black; not passing quality control) barcodes. **B)** Distribution of doublet scores for all snRNA-seq nuclei that passed initial Cell Ranger (10x Genomics) and Seurat^23^ quality control. **C)** Median genes detected per nucleus for each snRNA-seq dataset. **D)** Median UMIs detected per nucleus for each snRNA-seq dataset. **E)** Distribution of genes per nucleus on final snRNA-seq UMAP^59^. **F)** Distribution of UMIs per nucleus on final snRNA-seq UMAP^59^. **G)** Initial Seurat^23^ clustering result of snRNA-seq data showing 18 clusters, and dashed lines indicating final 12 major cell cluster annotations based on shared expression patterns (**H**). **H)** Differential gene expression heatmap showing top 10 differentially expressed genes for each initial cluster by Seurat^23^. Initial clusters were merged into major cell clusters based on shared gene expression patterns as indicated above the heatmap. **I, J)** UMAPs^59^ showing chamber-of-origin for nuclei included in the final (**I**) snRNA-seq and (**J**) snATAC-seq datasets.

**Supplemental Figure III:**
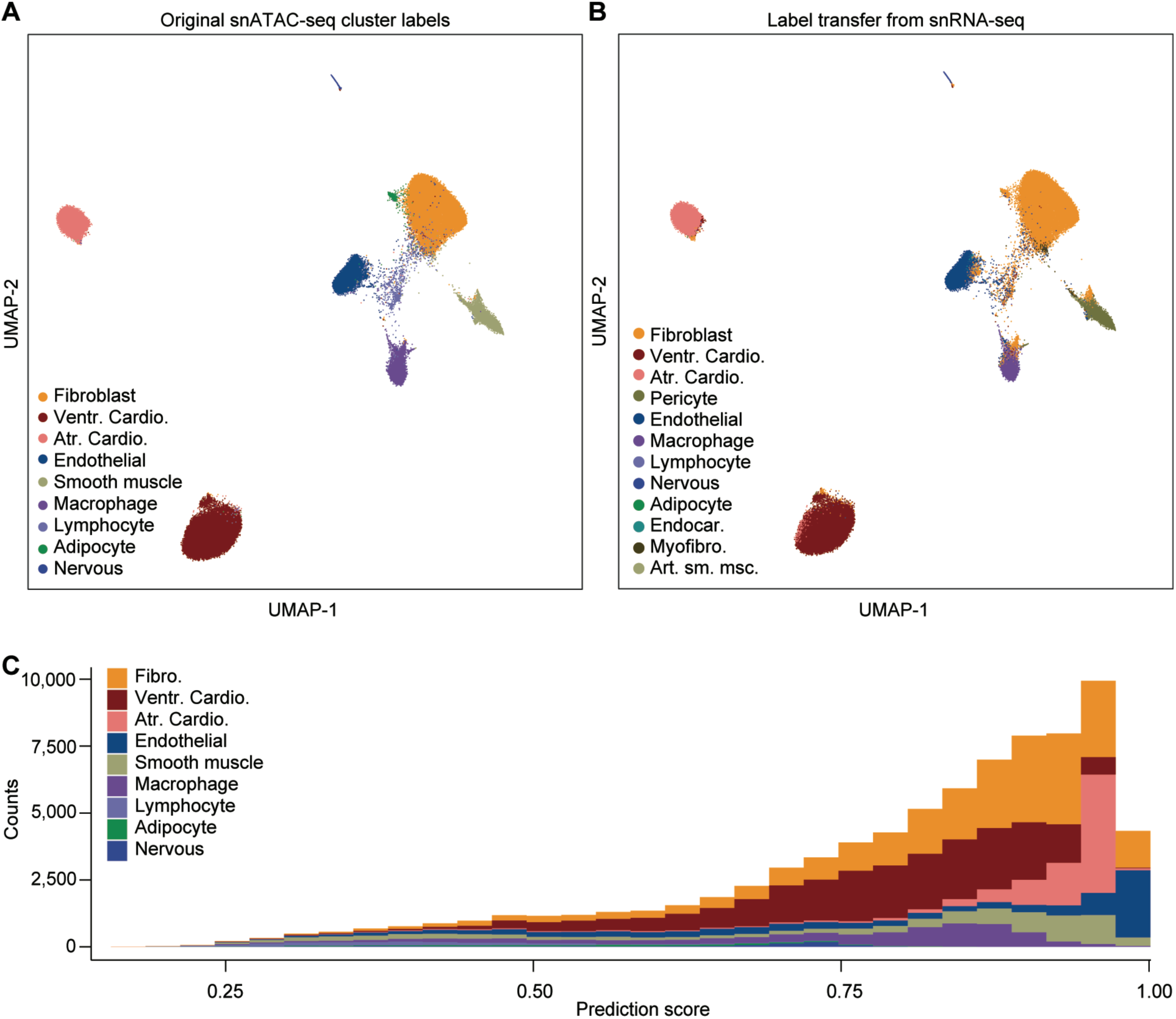
Integration of snRNA-seq and snATAC-seq datasets. **A, B)** Seurat^23^ was used to perform integration of chromatin accessibility and transcriptomes at the single cell level. **A)** UMAP^59^ showing nuclei colored based on original snATAC-seq cluster annotation (same as in Figure 1B). **B)** UMAP^59^ showing nuclei colored with cluster labels transferred from snRNA-seq. **C)** Histogram showing the prediction score distribution by original snATAC-seq cluster annotation. 93% of nuclei showed a prediction score >0.5 indicating a match between chromatin accessibility and transcriptome.

**Supplemental Figure IV:**
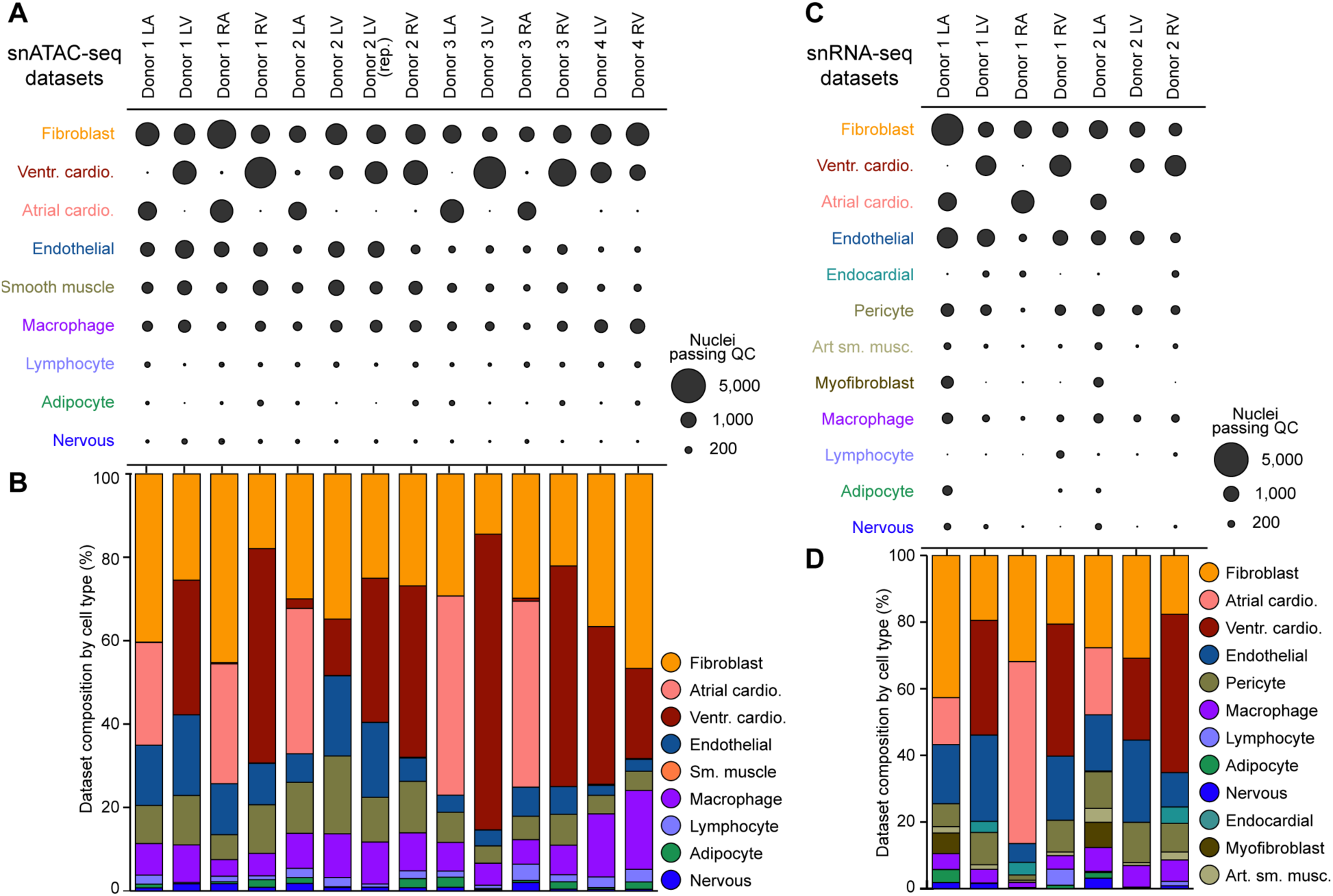
Cellular composition of snATAC-seq and snRNA-seq datasets. **A)** Dot plot showing number of nuclei passing quality control per cluster for each snATAC-seq dataset. **B)** Bar plot showing cell type composition of each snATAC-seq dataset as percentage of cell types. **C)** Dot plot showing number of nuclei passing quality control per cluster for each snRNA-seq dataset. **D)** Bar plot showing cell type composition of each snRNA-seq dataset as percentage of cell types.

**Supplemental Figure V:**
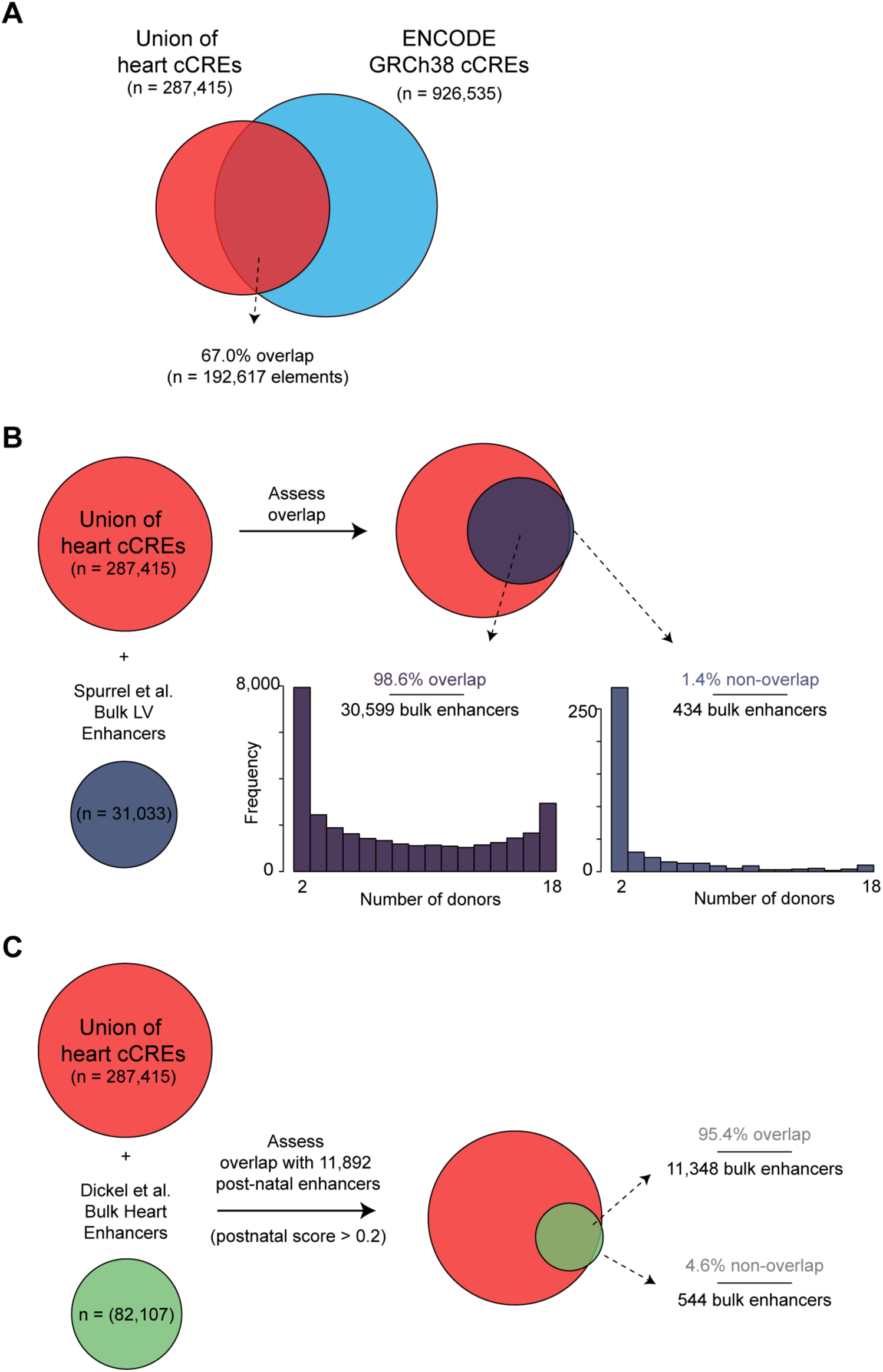
Overlap of union of heart candidate *cis* regulatory elements (cCREs) with several reference datasets. **A)** Overlap of the union of 287,415 heart cCREs from snATAC-seq with annotated cCREs in the human genome from the SCREEN database^26, 27^. **B)** Overlap of union with healthy left ventricular candidate enhancers from 18 human donors^14^. Arrows pointing from Venn diagram indicate number of overlapping (by at least one base pair) and non-overlapping genomic regions. Histograms display the number of donors harboring reported healthy heart enhancers (out of 18) for candidate enhancers that overlap union cCREs (left) and candidate enhancers that do not overlap union cCREs (right). **C)** Overlap of heart cCREs with post-natal heart candidate enhancers (reported post-natal score > 0.2) from a meta-analysis of epigenomic data from human and mouse heart tissues^12^. Venn diagrams are not to scale.

**Supplemental Figure VI:**
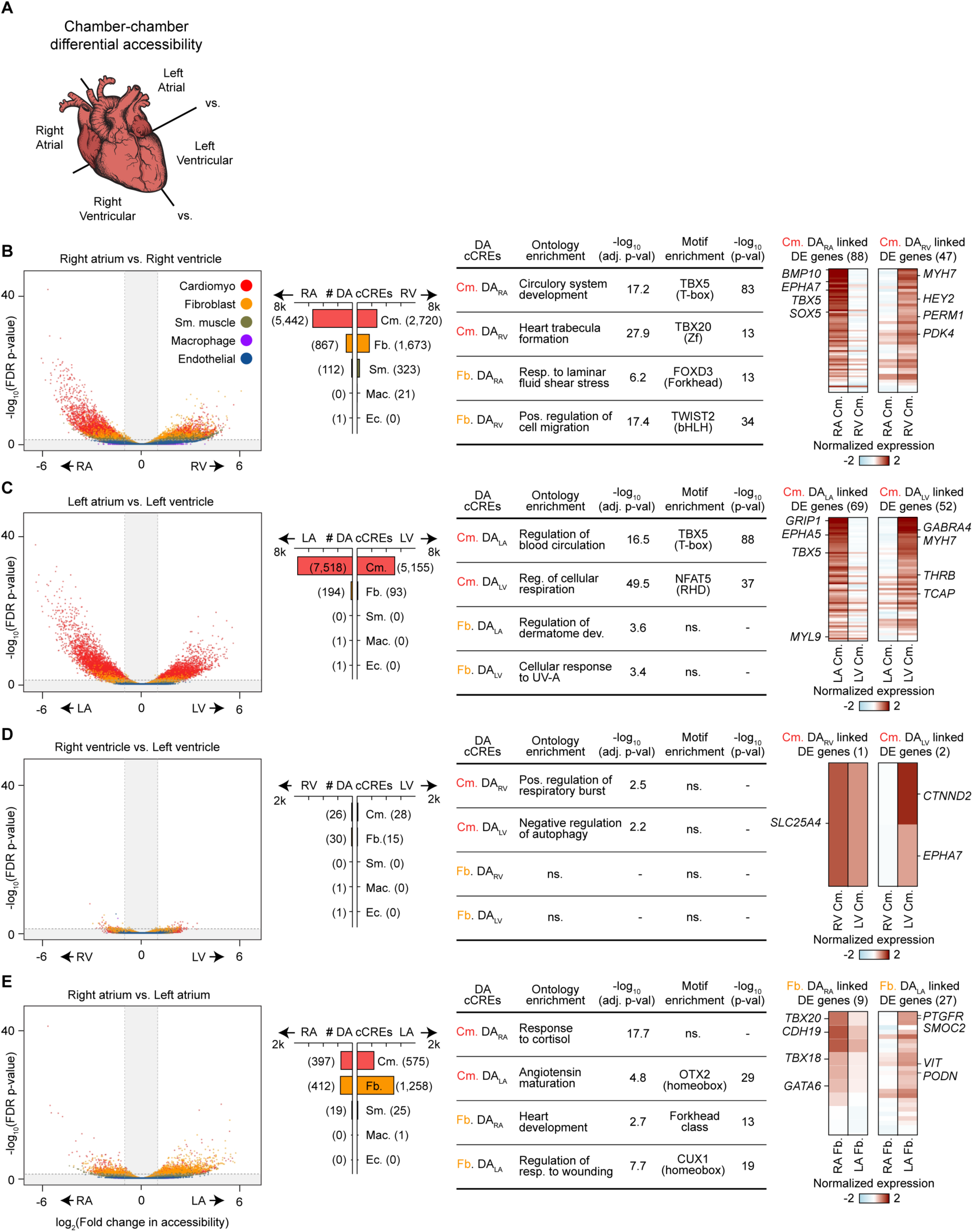
cCREs in cardiomyocytes and cardiac fibroblasts display chamber-dependent differences in accessibility. **A)** Scheme for comparison of major cell types across individual heart chambers to identify differential accessible (DA) cCREs. (**B-E**) Comparisons were performed between (**B**) right atrium (RA) and right ventricle (RV), (**C**) left atrium (LA) and left ventricle (LV), (**D**) right ventricle (RV) and left ventricle (LV) and (**E**) right atrium (RA) and left atrium (LA). For each comparison the following data are displayed. Left: Volcano plots showing identification of differentially accessible (DA) cCREs in each cell type between indicated chambers. cCREs with log_2_(fold change) > 1 and FDR < 0.05 after Benjamini-Hochberg correction (outside the shaded area) were considered DA. Each dot represents a cCRE and the color indicates the cell type. Second from the left: Bar plots showing number of DA cCREs per cell type. Number of DA cCREs listed in brackets. Second from the right: GREAT ontology analysis^28^ and transcription factor motif enrichment analysis result^30^ for the indicated DA cCREs. The best matches for selected *de novo* motifs (score > 0.7) are displayed. Statistical test for motif enrichment: hypergeometric test. P-values were not corrected for multiple testing. Ontology p-values were adjusted using Bonferroni correction. Right: Heatmaps showing normalized gene expression levels of differentially expressed genes linked to distal DA cCREs. Displayed are expression levels for putative target genes of distal DA cCREs for the cell type with most DA cCREs for the indicated chamber comparisons. Number of genes is shown in brackets. For lists of differentially expressed genes linked to distal DA cCREs for all comparisons in cardiomyocytes and fibroblasts see Supplemental Table XII (Cm. = cardiomyocyte, Fb. = fibroblast, ns. = no significant enrichment).

**Supplemental Figure VII:**
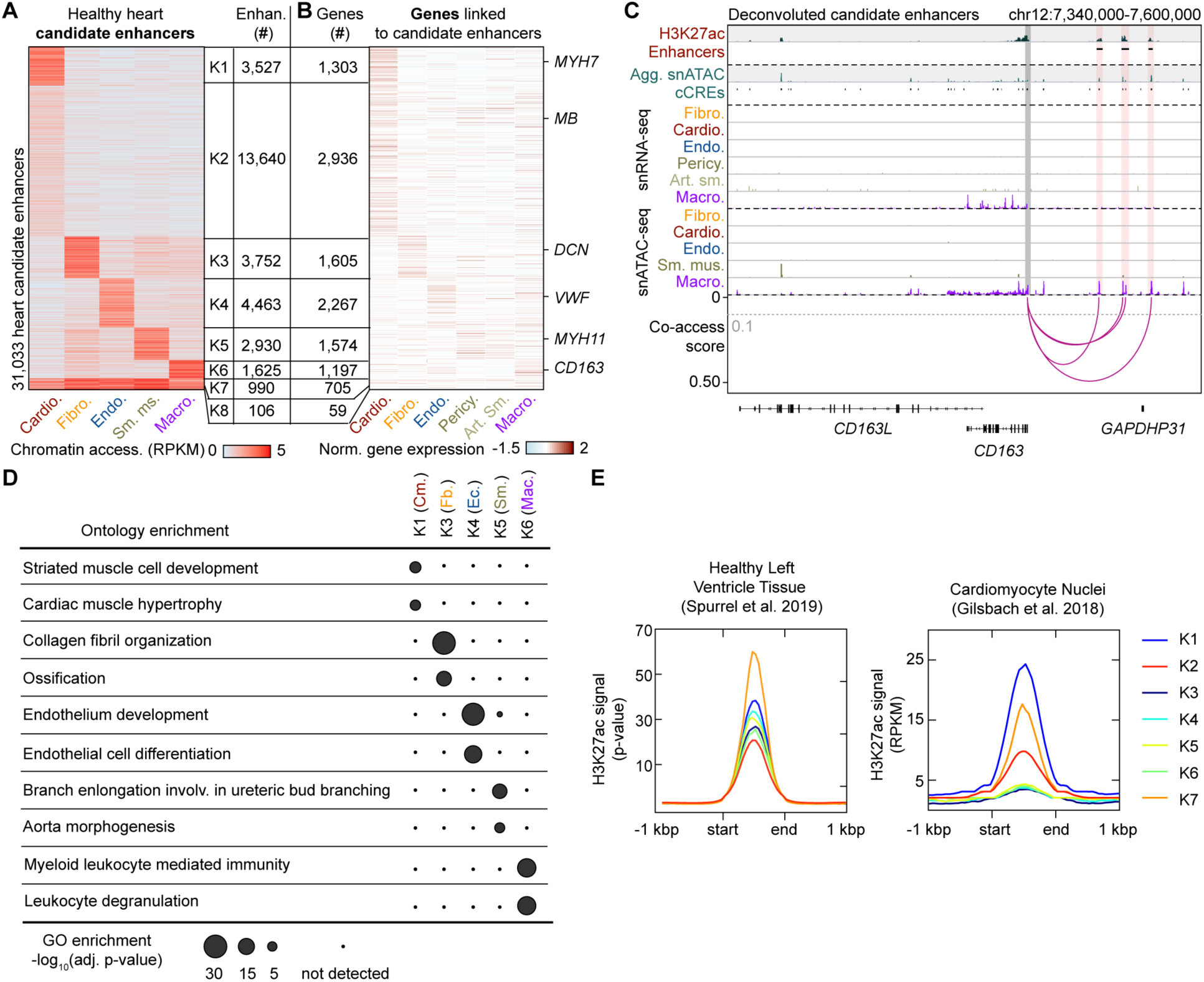
Deconvolution of candidate heart enhancers identified from bulk assays. **A)** H3K27ac peaks from bulk healthy heart tissue samples^14^ were deconvoluted into major cardiac cell types using cell type-resolved chromatin accessibility data. Heatmap displays cell type-resolved chromatin accessibility RPKM (reads per kilobase per million mapped reads) values from left ventricular snATAC-seq datasets. Candidate enhancers were grouped based on chromatin accessibility pattern across cell clusters using K-means. **B)** Heatmap displays cell-type resolved gene expression of putative enhancer target genes from left ventricular snRNA-seq datasets. **C)** Genome browser tracks^60^ of H3K27ac in left ventricle tissue and cell type-resolved gene expression (snRNA-seq) and chromatin accessibility (snATAC-seq) for several candidate heart enhancers (indicated by shaded red boxes) attributed to macrophages (K6 in panel **A**). The co-accessibility track shows linkages between the deconvoluted candidate enhancers and the promoter of *CD163* (cutoff > 0.1, grey dotted line). **D** GREAT analysis^28^ of deconvoluted candidate enhancers. Gene ontology enrichments are shown as Bonferroni-adjusted p-values. **E** Pileup tracks showing H3K27ac signal in bulk left ventricle datasets^14^ (left) and from purified cardiomyocyte nuclei^15^ (right) from non-failing (NF) hearts in distinct groups of enhancers which were either associated with a cell type (K1-6 in panel **A**) or broadly accessible across cell types (K7 in panel **A**). H3K27ac signal in cardiomyocyte nuclei data was highest in the cardiomyocyte-attributed candidate enhancers as well as the widely accessible candidate enhancers (K1,2,7), whereas signal strength in left ventricular tissue was highest in widely accessible enhancers and comparable between groups of cell type-specific candidate enhancers.

**Supplemental Figure VIII:**
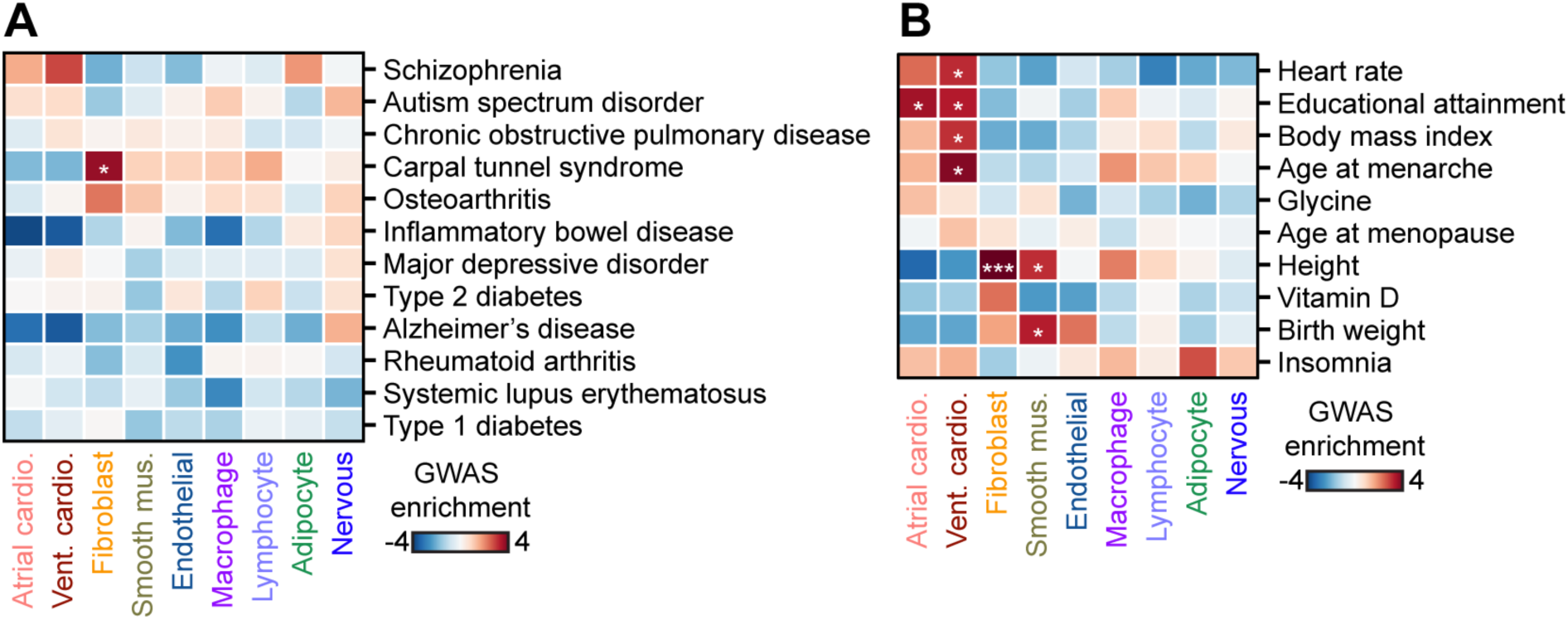
Risk variant enrichment analysis for non-cardiovascular diseases and non-disease traits. **A, B)** Enrichment of risk variants associated with (**A**) non-cardiovascular diseases and (**B**) non-disease traits from GWAS in cardiac cell type-resolved cCREs. Displayed are z-scores, and these scores were also used to compute one-sided p-values for enrichment that were corrected using the Benjamini Hochberg procedure for multiple testing (* = FDR < 0.05, *** = FDR < 0.001).

**Supplemental Figure IX:**
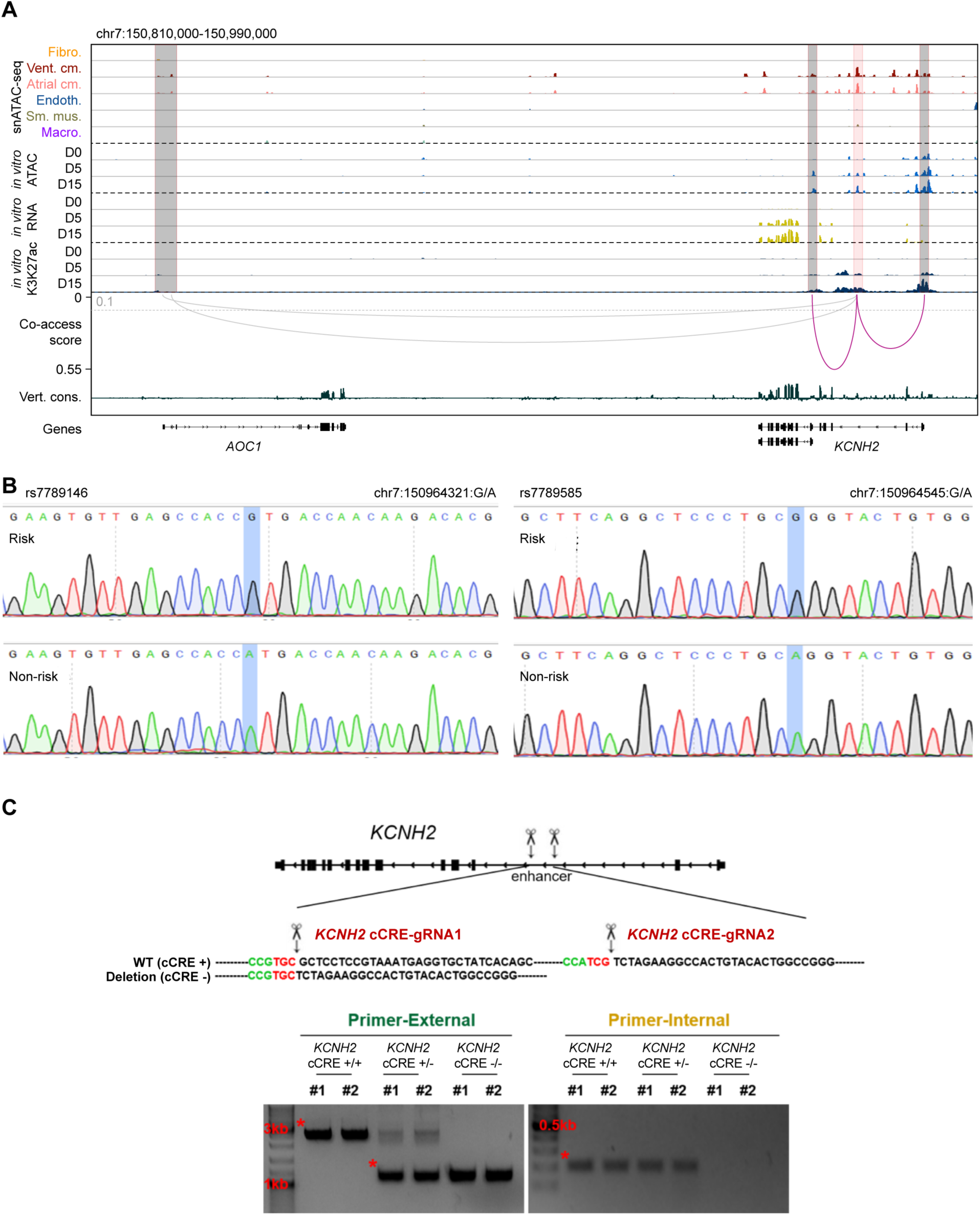
Validation of *KCNH2*-associated candidate enhancer. **A)** Genome browser tracks^60^ displaying cell type-resolved chromatin accessibility and co-accessibility from snATAC-seq as well as chromatin accessibility, H3K27ac signal, and gene expression during hPSC-cardiomyocyte differentiation. For illustration purposes, the co-accessibility track shows linkages between the AF variant-containing cCRE and annotated gene promoters (co-accessibility > 0.1, grey dotted line). The grey arc represents links to the promoter of *AOC1* which was not expressed. Figure 5C shows a zoom into this locus. **B)** Representative Sanger sequencing peak map at *KCNH2* intronic cCRE showing the risk allele for AF (top row, homozygous rs7789146-G / rs7789585-G) and the non-risk allele for AF (bottom row, homozygous rs7789146-A / rs7789585-A) used for luciferase assay. Blue highlighted regions indicate positions of variants. **C)** Schematic representation of the strategy for deletion of the *KCNH2* enhancer. The paired gRNAs (gRNA-1 and gRNA-2) were designed to target upstream and downstream of the *KCNH2* enhancer. Bottom panels show genomic DNA PCR verification of deletion in the H9-hTnnTZ-pGZ-D2 cell line. The red asterisk indicates specific bands.

## SUPPLEMENTAL TABLES

**Supplemental Table I.** Clinical metadata for heart samples.

**Supplemental Table II.** Quality control and cell type composition data for each snATAC dataset.

**Supplemental Table III.** Quality control, cell type composition, and integration with snATAC-seq results for snRNA-seq datasets.

**Supplemental Table IV.** snRNA-seq gene expression by major cluster and major cluster-specific genes. Included are genes that are expressed at higher (positive fold change) and lower levels (negative fold change) in a given cluster relative to the other clusters.

**Supplemental Table V.** Union of 287,415 cCREs in the cell types of the human heart.

**Supplemental Table VI.** List of 19,447 cell type-specific cCREs.

**Supplemental Table VII.** GREAT^28^ analysis for cell type-specific cCREs. Listed are biological processes with Bonferroni corrected p value <0.05.

**Supplemental Table VIII.** ChromVAR^29^ motif enrichment results in snATAC-seq cell clusters.

**Supplemental Table IX.** HOMER^30^ motif enrichment results for cell type-specific cCREs. Both *de novo* (p value < 10^-11^) and known motif (q value < 0.05) enrichments are reported.

**Supplemental Table X.** Differentially accessible cCREs between heart chambers.

**Supplemental Table XI.** Co-accessible cCRE pairs (score > 0.1) from Cicero^33^.

**Supplemental Table XII.** Lists of differentially accessible (DA) cCREs linked to differentially expressed genes.

**Supplemental Table XIII.** GREAT^28^ analysis for differentially accessible (DA) cCREs between heart chambers in cardiomocytes and fibroblasts. Listed are biological processes with Bonferroni corrected p value < 0.05.

**Supplemental Table XIV.** HOMER^30^ motif enrichments for differentially accessible (DA) cCREs between heart chambers in cardiomyocytes and fibroblasts. Both *de novo* (p value < 10^-11^) and known motif (q value < 0.05) enrichments are reported. CM: cardiomyocytes, FB: fibroblasts, LV: left ventricle, RV: right ventricle, LA: left atrium, RA: right atrium. For example, RA-vs-RV_CM-RV-DA_denovo denotes *de novo* motif enrichment in cardiomyocytes (CM) with higher accessibility (DA) in the right ventricle (RV) as compared to the right atrium (RA).

**Supplemental Table XV.** RPKM values and cluster membership for deconvoluted healthy and disease-associated candidate heart enhancers.

**Supplemental Table XVI.** GREAT^28^ analysis for distinct groups of deconvoluted candidate heart enhancers. Listed are biological processes with Bonferroni corrected p value < 0.05.

**Supplemental Table XVII.** HOMER^30^ motif enrichment results for distinct groups of deconvoluted candidate heart enhancers. Both *de novo* (p value < 10^-11^) and known motif (q value < 0.05) enrichments are reported.

**Supplemental Table XVIII.** Studies for non-cardiovascular disease and non-disease trait GWAS used for LD score regression.

**Supplemental Table XIX.** 38 fine mapped risk variants associated with atrial fibrillation within cardiomyocyte cCREs.

**Supplemental Table XX.** Primer sequences with indexes for snATAC-seq libraries.

**Supplemental Table XXI.** Primer sets used in qPCR assays.

## ONLINE METHODS

### Human Tissues

Adult human heart tissues were procured at the time of organ donation using an Institutional Review Board protocol (No. 101021) approved by the University of California, San Diego. Donated hearts were perfused with cold cardioplegia prior to cardiectomy and then explanted immediately into an ice-cold physiologic solution as we previously described^63^. Full-thickness samples from each chamber were obtained and epicardial fat rapidly removed before immediately flash freezing samples in liquid nitrogen. Samples were received from the United Network for Organ Sharing. Limited clinical data was obtained for each heart per approved Institutional Review Board protocol (Supplemental Table I). All samples were stored at −80°C until processing.

### Single nucleus ATAC-seq

Combinatorial barcoding single nucleus ATAC-seq was performed as described previously^17, 18, 22^ with slight modifications and using new sets of oligos for tagmentation and PCR (Supplemental Table XX). Nuclei were isolated in gentleMACS M-tubes (Miltenyi) on a gentleMACS Octo Dissociator (Miltenyi) using the “Protein_01_01” protocol in MACS buffer (5 mM CaCl_2_, 2 mM EDTA, 1X protease inhibitor (Roche, 05-892-970-001), 300 mM MgAc, 10 mM Tris-HCL pH 8, 0.6 mM DTT). Nuclei were pelleted with a swinging bucket centrifuge (500 x g, 5 min, 4°C; 5920R, Eppendorf) and resuspended in 1 mL Nuclear Permeabilization Buffer (1X PBS, 5% Bovine Serum Albumin, 0.2% IGEPAL CA-630 (Sigma), 1 mM DTT, 1X Protease inhibitor). Nuclei were rotated at 4 °C for 5 minutes before being pelleted again with a swinging bucket centrifuge (500 x g, 5 min, 4°C; 5920R, Eppendorf). After centrifugation, permeabilized nuclei were resuspended in 500 μL high salt tagmentation buffer (36.3 mM Tris-acetate (pH = 7.8), 72.6 mM potassium-acetate, 11 mM Mg-acetate, 17.6% DMF) and counted using a hemocytometer. Concentration was adjusted to 2,000 nuclei/9 μl, and 2,000 nuclei were dispensed into each well of a 96-well plate per sample (96 tagmentation wells/sample, samples were processed in batches of 2-4 samples). For tagmentation, 1 μL barcoded Tn5 transposomes (Supplemental Table XX) were added using a BenchSmart™ 96 (Mettler Toledo), mixed five times, and incubated for 60 min at 37 °C with shaking (500 rpm). To inhibit the Tn5 reaction, 10 μL of 40 mM EDTA (final 20mM) were added to each well with a BenchSmart™ 96 (Mettler Toledo) and the plate was incubated at 37 °C for 15 min with shaking (500 rpm). Next, 20 μL of 2x sort buffer (2 % BSA, 2 mM EDTA in PBS) were added using a BenchSmart™ 96 (Mettler Toledo). All wells were combined into a separate FACS tube for each sample, and stained with Draq7 at 1:150 dilution (Cell Signaling). Using a SH800 (Sony), 20 nuclei per sample were sorted per well into eight 96-well plates (total of 768 wells) containing 10.5 μL EB (25 pmol primer i7, 25 pmol primer i5, 200 ng BSA (Sigma)). During the sort, nuclei with 2-8 copies of DNA (2-8n) were included since cardiomyocyte nuclei in human hearts are often polyploid^15^. Preparation of sort plates and all downstream pipetting steps were performed on a Biomek i7 Automated Workstation (Beckman Coulter). After addition of 1 μL 0.2% SDS, samples were incubated at 55 °C for 7 min with shaking (500 rpm). 1 μL 12.5% Triton-X was added to each well to quench the SDS. Next, 12.5 μL NEBNext High-Fidelity 2× PCR Master Mix (NEB) were added and samples were PCR-amplified (72 °C 5 min, 98 °C 30 s, (98 °C 10 s, 63 °C 30 s, 72°C 60 s) × 12 cycles, held at 12 °C). After PCR, all wells were combined. Libraries were purified according to the MinElute PCR Purification Kit manual (Qiagen) using a vacuum manifold (QIAvac 24 plus, Qiagen) and size selection was performed with SPRISelect reagent (Beckmann Coulter, 0.55x and 1.5x). Libraries were purified one more time with SPRISelect reagent (Beckman Coulter, 1.5x). Libraries were quantified using a Qubit fluorimeter (Life technologies) and a nucleosomal pattern of fragment size distribution was verified using a Tapestation (High Sensitivity D1000, Agilent). Libraries were sequenced on a NextSeq500 sequencer (Illumina) using custom sequencing primers with following read lengths: 50 + 10 + 12 + 50 (Read1 + Index1 + Index2 + Read2). Primer and index sequences are listed in Supplemental Table XX.

### Single nucleus RNA-seq

Nuclei were isolated from heart tissue using a gentleMACS (Miltenyi) dissociator. ∼40 mg of frozen heart tissue was suspended in 2 ml of MACS dissociation buffer (5 mM CaCl2 (G-Biosciences, R040), 2 mM EDTA (Invitrogen, 15575-038), 1X protease inhibitor (Roche, 05-892-970-001), 3 mM MgAc (Grow Cells, MRGF-B40), 10 mM Tris-HCl pH 8 (Invitrogen, 15568-075), 0.6 mM DTT (Sigma-Aldrich, D9779), and 0.2 U/μL of RNase inhibitor (Promega, N251B) in water (Corning, 46-000-CV)) and placed on wet ice. Next, samples were homogenized using gentleMACS dissociator (Miltenyi) with gentleMACS M tubes (Miltenyi, 130-096-335)) and the “Protein_01_01” protocol. Suspension was filtered through a 30 μM CellTrics filter (Sysmex, 04-0042-2316). M tube and filter were washed with 3 mL of MACS dissociation buffer and combined with the suspension. Suspension was centrifuged in a swinging bucket centrifuge (Eppendorf, 5920R) at 500 g for 5 minutes (4°C, ramp speed 3/3). Supernatant was carefully removed and pellet was resuspended in 500 μL of nuclei permeabilization buffer (0.1% Triton X-100 (Sigma-Aldrich, T8787), 1X protease inhibitor (Roche, 05-892-970-001), 1 mM DTT (Sigma-Aldrich, D9779), 0.2 U/μL RNase inhibitor (Promega, N251B), and 2% BSA (Sigma-Aldrich, SRE0036) in PBS). Sample was incubated on a rotator for 5 minutes at 4°C and then centrifuged at 500 g for 5 minutes (Eppendorf, 5920R; 4°C, ramp speed 3/3). Supernatant was removed and pellet was resuspended in 600-1000 μl of sort buffer (1 mM EDTA and 0.2 U/μL RNase inhibitor in 2% BSA (Sigma-Aldrich, SRE0036) in PBS) and stained with DRAQ7 (1:100, Cell Signaling, 7406). 75,000 nuclei were sorted using a SH800 sorter (Sony) into 50 μL of collection buffer (1 U/ μL RNase inhibitor, 5% BSA (Sigma-Aldrich, SRE0036) in PBS); Sorted nuclei were then centrifuged at 1000 g for 15 minutes (Eppendorf, 5920R; 4°C, ramp speed 3/3) and supernatant was removed. Nuclei were resuspended in 18-25 ul of reaction buffer (0.2 U/μL RNase inhibitor, 1% BSA (Sigma-Aldrich, SRE0036) in PBS) and counted using a hemocytometer. 12,000 nuclei were loaded onto a Chromium controller (10x Genomics). Libraries were generated using the Chromium Single Cell 3′ Library Construction Kit v3 (10x Genomics, 1000078) according to manufacturer specifications. cDNA was amplified for 12 PCR cycles. SPRISelect reagent (Beckman Coulter) was used for size selection and clean-up steps. Final library concentration was assessed by Qubit dsDNA HS Assay Kit (Thermo-Fischer Scientific) and fragment size was checked using Tapestation High Sensitivity D1000 (Agilent) to ensure that fragment sizes were distributed normally around 500 bp. Libraries were sequenced using a NextSeq500 or HiSeq4000 (Illumina) using these read lengths: Read 1: 28 cycles, Read 2: 91 cycles, Index 1: 8 cycles.

### Human pluripotent stem cell culture

An engineered H9-hTnnTZ-pGZ-D2 human pluripotent stem cell transgenic reporter line was purchased from WiCell and maintained on Geltrex (Gibco) pre-coated tissue culture plates in E8 medium^64^ containing DMEM/F12, L-ascorbic acid-2-phosphate magnesium (64 mg/L), sodium selenium (14 μg/L), FGF2 (100 μg/L), insulin (19.4 mg/L), NaHCO3 (543 mg/L) transferrin (10.7 mg/L), and TGFβ1(2 μg/L). Cells were passaged every 3 to 5 days upon reaching ∼80% confluency. For single cell passaging experiments, cells were incubated with pre-warmed TrypLE™ Select Enzyme, no phenol red (1 mL per well of a 6-well plate) for 2-3 minutes in a 37°C, 5% CO2 incubator. Following incubation, cells were triturated to create a single cell suspension and cultured in E8 Medium supplied with Rock inhibitor^65^ for 18-24 hours post-split, followed by daily feeding with E8 medium.

### *In vitro* cardiomyocyte differentiation

The H9-hTnnTZ-pGZ-D2 cell line was differentiated into beating cardiomyocytes utilizing a previously reported Wnt-based monolayer differentiation protocol^66^. Briefly, the H9-hTnnTZ-pGZ-D2 cell line was cultured in E8 medium for 3-10 passages. Prior to differentiation, human pluripotent stem cells were seeded at a density of 350,000-400,000 cells per well of a 12-well plate and cultured for two days. For direct differentiation, cells were treated with 10 μM CHIR99021 (Fisher, #442350) in RPMI/B-27 without insulin. Fresh RPMI/B-27 without insulin media was replaced at post 24hr and cells were then cultured two days. At day 3, cells were treated with 5 μM IWP2 (TOCRIS, #353310) in conditional medium and RPMI/B-27 without insulin 1:1 mix medium for another two days. At day 5, cells were exposed to fresh RPMI/B-27 without insulin media again for two days. Then, fresh RPMI/B-27 with insulin media was used and replenished every two days. Contracting cardiomyocytes were usually observed at day 7-8. D25 in vitro cardiomyocytes were purified utilizing PSC-derived cardiomyocyte isolation kit, human (Miltenyi Biotec, 130-110-188) and used for Real-time quantitative PCR (RT-qPCR).

### Luciferase reporter assay

A genomic region harboring the *KCNH2* intronic enhancer (containing the risk allele: homozygous rs7789146-G / rs7789585-G) was amplified by nested-PCR using genomic DNA of H9-hTnnTZ-pGZ-D2 transgenic cells as a template and cloned into pGL4.23 [luc2/minP] (Promega, Cat#E8411) luciferase reporter vector. Synthetic DNA containing the *KCNH2* intronic enhancer with the non-risk allele (homozygous rs7789146-A / rs7789585-A) was purchased from integrated DNA technologies and cloned into pGL4.23 [luc2/minP] luciferase vector. One day prior to transfection, 3×10^5^ of D15 *in vitro* differentiated cardiomyocytes were plated in a Geltrex-coated 24-well plate. Cardiomyocytes were transfected with 500 ng of pGL4.23 plasmid (either empty, *KCNH2* enhancer with G/G allele, or A/A allele) and 10 ng TK:Renilla-luc as internal control using Lipofectamine Stem Transfection Reagent (Invitrogen, #STEM00003). Media was replaced with fresh media at 24 hrs post-transfection. At 72 hrs post-transfection, media was removed and the cells were washed with PBS. Luminescence was measured using a Dual-Luciferase Reporter Assay System (Promega, #E2920) according to the manufacturer’s protocol.

### CRISPR mediated genome editing experiments

To interrogate the functional significance of the atrial fibrillation-associated risk variant-containing cCRE at the *KCNH2* locus, the cCRE sequence was genetically deleted in H9-hTnnTZ-pGZ-D2 transgenic hPSCs using an efficient CRISPR/Cas9-mediated knockout system^49, 67^. Two adjacent gRNAs (KCNH2-enh gRNA-1, CTCATTTACGGAGGAGCGCA; KCNH2-enh gRNA-2, TACAGTGGCCTTCTAGACGA) targeting the cCRE were designed using a web-based software tool CRISPOR^68^, based on targeting region of interest and minimizing potential off-target effects. The identified gRNAs were then synthesized in vitro using the GeneArt Precision gRNA Synthesis kit (Invitrogen) according to the manufacturer’s protocol. One day prior to transfection, 1.5×10^5^ H9-hTnnTZ-pGZ-D2 hPSCs were seeded in 12-well plates. A pair of RNP complexes containing 1.2 μg of Cas9 protein (NEB) and 400 ng of in vitro transcribed gRNA were then transfected^69, 70^ using Lipofectamine stem transfection reagent (Invitrogen). 72 hours after the transfection, cells were diluted and clonally expanded another 7 days. Colonies were picked and lysates were prepared after the first passage for genotyping^71^ (KCNH2-enh extended forward primer, ACACCTTACTTTGGGTGAGAAG; KCNH2-enh extended reverse primer, AGACAGAGCACAGACCTAGAA; KCNH2-enh internal forward primer, GCTGTGCAGTGTCAGGTTAT; KCNH2-enh internal reverse primer, TCTCCCTCCTTCTCTCTCATTC). After confirmation of genome-edited clones by Sanger sequencing, two transfected WT clones, two heterozygote clones, and two homozygote clones were selected for further functional analysis.

### RT-qPCR

Total RNA was isolated from the cells using TRIzol reagent (Invitrogen). 1 μg of total RNA was reverse transcribed using the iScript Reverse Transcription Supermix kit (Bio-Rad) for RT-qPCR. RT-qPCR was performed using PowerUP^TM^ SYBR^TM^ Green Master Mix (Applied Biosystems) in the CFX Connect Real-Time System (Bio-Rad). The results were normalized to the *TBP* gene. The primers used for RT-qPCR are listed in Supplemental Table XXI.

### Electrophysiology of cardiomyocytes

Both WT and *KCNH2* enhancer knockout D15 *in vitro* cardiomyocytes were purified using the PSC-derived cardiomyocyte isolation kit, human (Miltenyi Biotec, 130-110-188) and cultured for another 10-20 days in a low density prior to electrophysiological measurements. The single-pipette, whole-cell patch current-clamp technique was used for recordings. Action potentials were recorded with a patch clamp amplifier (Axopatch 200B, Axon) and experiments were performed at a temperature of 35 ± 0.5 °C. Current-clamp command pulses were generated by a digital-to-analog converter (DigiData 1440, Axon) which was controlled by the pCLAMP software (10.3, Axon). Pipettes (resistance 3-5 MΩ) were pulled using a micropipette puller (Model P-87, Sutter Instrument Co.). Several minutes after seal formation, the membrane was ruptured by gentle suction to establish the whole-cell configuration for voltage clamping. Subsequently, the amplifier was switched to the current-clamp mode. Cells were paced with 1 Hz, injected current stimuli from 3 to15 nA for 5 ms duration. Cells were superfused with extracellular solution containing (in mM): 140 NaCl, 5.4 KCl, 1.8 CaCl_2_, 1.0 MgCl_2_, 5.5 glucose and 5.0 HEPES (pH 7.4 adjusted with NaOH). Pipette solution contained (in mM): 120 K-gluconate, 10 KCl, 5 NaCl, 10 HEPES, 5 Phosphocreatine, 5 ATP-Mg_2_ and Amphotericin 0.44 μM (pH 7.2 adjusted with KOH).

## DATA ANALYSIS

### Demultiplexing of snATAC-seq reads

For each sequenced snATAC-Seq library, we obtained four FASTQ files, two for paired-end DNA reads as well as the combinatorial indexes for i5 (768 different PCR indices) and T7 (96 different tagmentation indices; Supplemental Table XX). We selected all reads with <= 2 mistakes per individual index (Hamming distance between each pair of indices is 4) and subsequently integrated the full barcode at the beginning of the read name in the demultiplexed FASTQ files (https://gitlab.com/Grouumf/ATACdemultiplex/).

### Filtering of snATAC-seq profiles by TSS enrichment and unique fragments

TSS (transcriptional start site) positions were obtained from the GENCODE database v31^61^. Tn5-corrected insertions were aggregated ± 2000 bp around each TSS genome wide. Then, this profile was normalized to the mean accessibility ± (1900 to 2000) bp from the TSS and smoothed every 11 bp. The maximum value of the smoothed profile was taken as the TSS enrichment. We selected all nuclei that had at least 1,000 unique fragments and a TSS enrichment of at least 7 for all data sets.

### Clustering strategy for snATAC-seq datasets

We utilized two rounds of clustering analysis to identify clusters. The first round of clustering analysis was performed on individual samples. We divided the genome into 5 kbp consecutive bins and then scored each nucleus for any insertions in these bins, generating a bin-by-cell binary matrix for each sample. We filtered out those bins that are generally accessible in all nuclei for each sample using z-score threshold 1.65. Based on the filtered matrix, we then carried out dimensionality reduction followed by graph-based clustering to identify cell clusters. We called peaks using MACS2^25^ for each cluster using the aggregated profile of accessibility and then merged the peaks from all clusters to generate a union peak list. Based on the peak list, we generated a cell-by-peak count matrix and used Scrublet^62^ to remove potential doublets with default parameters. Doublet scores returned by Scrublet^62^ were then used to fit a two-component Gaussian mixture model using the *BayesianGaussianMixture* function from the python package *scikit-learn*^72^. Nuclei in the component with the larger mean doublet score were removed from downstream analysis since they likely reflected doublets.

Next, to carry out the second round of clustering analysis, we merged peaks called from all samples to form a reference peak list. We generated a binary cell-by-peak matrix using nuclei from all samples and again performed the dimensionality reduction followed by graph-based clustering to obtain the final cell clusters across the entire dataset.

### Dimensionality reduction and batch correction of snATAC-seq data

For processing of snATAC-seq data we adapted our previously published method, SnapATAC^22^. To reduce the dimensionality of the peak by cell count matrix, SnapATAC utilizes spectral embedding for dimensionality reduction. To further increase the performance and scalability of spectral embedding, we applied the Nyström method^73^ to enable handling of large datasets. Specifically, we first randomly sampled 35,000 nuclei as training data. We then computed the Jaccard index between each pair of cells in the training set and constructed the similarity matrix S. We computed the matrix *P* = *D*^−1^*S* S, where D is the diagonal matrix such that *D_ii_* = ∑*_j_ S_ij_*. The eigendecomposition was performed on P and the eigenvector with eigenvalue 1 was discarded. From the rest of the eigenvectors, we took k of them corresponding to the largest eigenvalues as the spectral embedding of the training data. We utilized the Nyström method^73^ to extend the embedding to the data outside the training set. Given a set of unseen samples, we computed the similarity matrix S’ between the new samples and the training set. The embedding of the new samples is given by ′ = *S*′*UΛ*^−1^, where U and Λ are the eigenvectors and eigenvalues of P obtained in the previous step.

To correct for donor/batch specific effects, after dimensionality reduction we performed cell grouping on individual samples using k-mean clustering with k equal to 20. We then constructed k-NN graphs for each sample and used the MNN correction method to identify mutual nearest neighbors^74^.These mutual nearest neighbors were used as the anchors to match the cells between different samples and correct for donor/batch effects as described previously^74^.

### Clustering of snATAC-seq data

We constructed the k-nearest neighbor graph (k-NNG) using low-dimensional embedding of the nuclei with k equal to 50. We then applied the Leiden algorithm^75^ with constant Potts model (CPM) to find communities in the k-NNG corresponding to the cell clusters. The Leiden algorithm can be configured to use different quality functions. The modularity model is a popular choice but it is hampered by the resolution-limit, particularly when the network is large^76^. Therefore, we used the modularity model only in the first round of clustering analysis to identify initial clusters. In the final round of clustering, we chose the constant Potts model as the quality function since it is resolution-limit-free and is better suited for identifying rare populations in a large dataset^76^. Nuclei from two small clusters (280 and 254 nuclei) with low reproducibility and stability were discarded from downstream analysis. 34 nuclei that formed clusters of 1 and 2 nuclei were discarded as well.

### Processing and clustering analysis of snRNA-seq datasets

Raw sequencing data was demultiplexed and preprocessed using the Cell Ranger software package v3.0.2 (10x Genomics). Raw sequencing files were first converted from Illumina BCL files to FASTQ files using *cellranger mkfastq*. Demultiplexed FASTQs were aligned to the GRCh38 reference genome (10x Genomics), and reads for exonic and intronic reads mapping to protein coding genes, long non-coding RNA, antisense RNA, and pseudogenes were used to generate a counts matrix using *cellranger count*; expect-cells parameter was set to 5,000. A separate counts matrix for each sample was also generated using only reads mapped to intronic regions.

Next, exon + intron count matrices for individual datasets were processed using the Seurat v3.1.4 R package^23^ (https://satijalab.org/seurat/) to assess dataset quality. Features represented in at least 3 cells and barcodes with between 500 and 4,000 genes were used for downstream processing; additionally, barcodes with mitochondrial read percentages greater than 5% were removed. Counts were log-normalized and scaled by a factor of 10,000 using *NormalizeData*. To identify variable genes, *FindVariableFeatures* was run with default parameters except for nfeatures = 3000 to return the top 3,000 variable genes. All genes were then scaled using *ScaleData*, which transforms the expression values for downstream analysis. Next, principal component analysis was performed using *RunPCA* with default parameters and the top 3,000 variable features as input. The first 20 principal components were used to run clustering using *FindNeighbors* and *FindClusters* (parameter res = 0.4). To generate UMAP coordinates *RunUMAP* was run using the first 20 principal components and with parameters umap.method = “umap-learn”, and metric = “correlation”. Doublet scores (pANN) were generated for cell barcodes using DoubletFinder ^77^ (https://github.com/chris-mcginnis-ucsf/DoubletFinder) using the parameters pN =0.15 and pK = 0.005; the anticipated collision rate was set by specifying 2% collisions per thousand nuclei for individual datasets.

Individual datasets were merged together using the *merge* function in Seurat to combine the count matrices and designate unique barcodes. Cell barcodes with pANN scores greater than 0 were removed from downstream analysis. Metadata was also encoded for each barcode, and the merged dataset was processed in a similar manner as described above; clusters were identified using *FindNeighbors* and *FindClusters* (res = 0.8). To generate the UMAP coordinates, the first 14 principal components were used in *RunUMAP*; the UMAP algorithm for Seurat v3.1.4 uses the uwot R-package, and that setting was used to generate the coordinates here. To regress out donor specific effects, the Harmony R package (https://github.com/immunogenomics/harmony)^78^ was used, and the recomputed principal components were used to re-cluster the cells and rerun UMAP using the above parameters. For downstream analysis and comparison to snATAC-seq data we combined ventricular cardiomyocyte clusters, atrial cardiomyocyte clusters, fibroblast clusters, and endothelial cell clusters manually based on shared gene expression patterns (Fig S2G, H). Cluster-specific genes in the all-transcripts dataset were identified in a global differential gene expression test using *FindAllMarkers* with parameters logFC = 0.25, min.pct = 0.25, and only.pos = FALSE.

### Integration of snRNA-seq and snATAC-seq data

The snRNA-seq and snATAC-seq datasets were used to perform label transfer from the RNA cells onto the snATAC-seq dataset using the Seurat v3.1.4 R package (https://satijalab.org/seurat/)^23^. Gene activity scores were calculated using chromatin accessibility in regions from the promoter up to 2kb upstream for each ATAC nucleus. Activity scores were log-normalized and scaled using *NormalizeData* and *ScaleData*. To compare the snRNA and snATAC datasets and identify anchors, *FindTransferAnchors* was run considering the top 3,000 variable features from the snRNA-seq dataset. Anchor pairs were used to assign RNA-seq labels to the snATAC-seq cells using *TransferData*, with the weight.reduction parameter set to the principal components used in snATAC-seq clustering. The efficacy of integration was assessed by examining the distribution of the maximum prediction scores output by *TransferData* and the distribution of annotated snATAC-seq identities to the corresponding predicted label.

### Creation of a consensus list of heart candidate *cis* regulatory elements

MACS2 (v2.1.2)^25^ was used to identify accessible chromatin sites for each cluster with the following parameters: *-q 0.01 --nomodel --shift -100 --extsize 200 -g 2789775646 --call-summits --keepdup-all.* Estimated genome size was determined to be 2789775646 bp and was indicated by the *-g* parameter. We next filtered out peaks overlapping with the ENCODE blacklist^79^ (hg38, https://github.com/Boyle-Lab/Blacklist/).

To generate the union of heart cCREs, we merged the blacklist-filtered peaks obtained for each cluster using the BEDtools merge command with default settings (v2.25.0) ^80^.

### Computing relative accessibility scores for candidate *cis* regulatory elements

To correct biases arising from differential read depth among cells and cell types, we derived a procedure that normalizes chromatin accessibility at cCREs identified by MACS2 peak calling (v2.1.2)^25^. We define the set of accessible loci by *L* and we define a peak *p* as a subset of related loci *l* from *L.* Let *a_l_* be the accessibility of accessible locus *l* and *P* the set of non-overlapping peaks used to define the loci. For a given cell type *S_i_* ∈ *S*, we computed the median *med_j_* number of reads sequenced per cells. For each feature *p_j_* ∈ *P*, we computed *m_ij_* the average number of reads sequenced from *S_i_* and overlapping *p_j_*. We then defined the activity *a_ij_* of loci *p_i_* in *S_j_* as 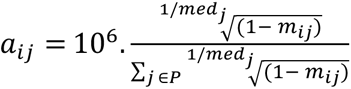. We then define the relative accessibility score (RAS) 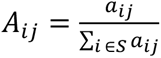.

### K-means clustering of candidate *cis* regulatory elements

We clustered the union of 287,415 candidate *cis* regulatory elements (cCREs) using a K-means clustering procedure. We first created a sparse cell x peak matrix that was transformed into a RAS-normalized cell type x peak matrix. We then performed K-means on the normalized matrix with *K* from 2 to 12 and computed the Davies-Bouldin (DB) index for each K^81^. Let 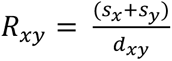 with *s_x_* the average distance of each cell of cluster *x* and *d_xy_* the distance between the centroids of clusters *x* and *y*. The Davies-Bouldin index is defined as 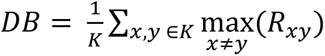. We selected K = 9 since it resulted in the lowest DB index which indicates the best partition. We used the python library scikit-learn^72^ to compute the K-means algorithm and the DB index^81^.

### Cell type annotation

We annotated snATAC-seq and snRNA-seq clusters based on chromatin accessibility at promoter regions or expression of known lineage marker genes, respectively. We annotated atrial and ventricular cardiomyocytes based on differential chromatin accessibility and gene expression at *NPPA, MYH6*, *KCNJ3*, *MYL7*, *MYH7, HEY2, MYL2* and other reported markers of atrial and ventricular cardiomyocytes^82–84^. We used, for example, the gene *DCN* to annotate cardiac fibroblasts^85^; *VWF* and *EGFL7* for endothelial cells^86, 87^; *GJA4* and *TAGLN* for smooth muscle cells^88, 89^; *CD163* and *MS4A6A* for macrophages^90, 91^; *IL7R* and *THEMIS* for lymphocytes^92, 93^; *ADIPOQ* and *CIDEA* for adipocytes^94, 95^; *NRXN3* and *GPM6B* for a cluster of nervous cells with neuronal and Schwann-like gene expression and chromatin accessibility signatures^9, 10, 96^. From snRNA-seq, we identified a population of endothelial-like cells with specific expression of endocardial cell markers *NRG3* and *NPR3*^97, 98^. We also identified subtypes of mesenchymal cells that included myofibroblasts with characteristic expression of embryonic smooth muscle actin *MYH10*^99, 100^ as well as arterial smooth muscle cells with preferential expression of *ACTA2* and *TAGLN* relative to a larger cluster of pericytes^101^ (Supplemental Table IV). snRNA-seq annotations were consistent with recent single cell transcriptomic analyses of adult human heart tissue^9, 10^.

### Identification of cell type-specific candidate *cis* regulatory elements

We used *edgeR* (version 3.24) in R^102^ to identify cell type-specific cCREs. For each cCRE, accessibility within a cell type was compared to average accessibility in all other clusters. For each cell type, we created a count table for each cCRE using the following strategy: each sample was described with a donor and a chamber ID. For each sample ID we reported read count within 1) the cell type and 2) the rest of the cell types in aggregate. We used this count matrix as input for edgeR analysis^102^. We performed a likelihood ratio test and considered peaks with FDR < 0.01 after Benjamini-Hochberg correction and log_2_(fold Change) > 0 as cell type-specific.

### Co-accessibility analysis using Cicero

We used the R package Cicero^33^ to infer co-accessible chromatin loci. For each chromosome, we used as input the corresponding peaks from our 287,415 cCRE union set and the coordinates of the snATAC-seq UMAP^59^. We randomly subsampled 15,000 cells from our aggregate snATAC-seq dataset to construct input matrices for Cicero analysis. We used +/−250 kbp as cutoff for co-accessibility interactions. All other settings were default.

### Correlation of gene expression and promoter accessibility

We defined promoter regions as transcriptional start sites (TSS) +/−2 kbp. Transcriptional start sites were extracted from annotation files from GENCODE release 33^61^. We identified promoter-overlapping peaks using BEDtools^80^ and a custom script (see **Code availability**). For each overlapping pair (peak, promoter) identified, we kept only the open chromatin site closest to the TSS in order to obtain a 1:1 correspondence between genes and open chromatin peaks. We then used the relative accessibility score (RAS) and the cluster-scaled FPKM gene expression score to create feature x cell type matrices for RNA-seq and ATAC-seq datasets. We then used these matrices to create heatmaps and to perform ATAC-seq/RNA-seq cluster correlation analysis using the Pearson similarity metric. For each cell type, we computed the Pearson correlation score between the RAS vector of the 7,081 promoters and the scaled FPKM vector of the corresponding 7,081 genes identified via the 1:1 correspondence method described above.

### Differential accessibility between cell types by chamber

Between-heart chamber differential accessibility analysis was performed for five cell types from our aggregated single nuclear ATAC-seq dataset. We considered only cell types which had a representation of at least 50 nuclei per dataset and at least 300 nuclei across each tested condition. The cell types that met these inclusion criteria included cardiomyocytes, fibroblasts, endothelial cells, smooth muscle cells, and macrophages. Within each cell type, a generalized linear model framework was employed using the R package edgeR^102^. All fragments for a given cell type were aggregated in the .bed format. MACS2^25^ was used to call peaks on the aggregate .bed file for each cell type with the parameters specified above. *NarrowPeak* output bed files were used for differential accessibility testing. The aggregate .bed file for each cell type was then partitioned based on dataset of origin using nuclear barcodes. The ‘*coverage*’ option of the BEDtools package^80^ was applied with default settings to count the total number of chromatin fragments from each dataset overlapping *narrowPeaks* called on the aggregate .bed file for the corresponding cell type. This yielded a raw count matrix in the format of single nuclear ATAC-seq datasets (columns) by *narrowPeaks* (rows) for each cell type. The raw count matrix was used as input for edgeR analysis. To filter low-coverage peaks from our analysis, we used the ‘*filterByExpr*’ command within edgeR with default settings. We applied an average prior count of one during fitting of the generalized linear model in order to avoid inflated fold changes in instances for which peaks lacked coverage for one but not both tested conditions. We modelled chromatin accessibility at each peak as a function of heart chamber (group) with sex as a covariate. The generalized linear model was expressed as follows in edgeR notation:

~~~
--
design <- model.matrix(∼sex+group)
y <- estimateDisp(y, design, prior.count = 1)
glmFit(y, design)
--
~~~

Significance was tested using a likelihood ratio test. To account for testing multiple hypotheses, a Benjamini-Hochberg significance correction was applied for all cCREs tested within each considered cell type. Any cCRE with an absolute log_2_(fold change) > 1 and an FDR-corrected p value < 0.05 was considered significant.

### Gene expression analysis of genes co-accessible with DA candidate *cis* regulatory elements

To compare the expression of genes co-accessible with heart chamber-dependent distal DA cCREs (outside +/− 2 kb of TSS) in cardiomyocytes and fibroblasts, we performed differential expression testing for all genes between indicated heart chambers using Wilcoxon rank sum test in Seurat^23^. Genes with an absolute Fold Change > 1.5 and an FDR-adjusted P value < 0.05 were considered differentially expressed. We then tested resulting genes for co-accessibility^33^ with distal DA cCREs at a co-accessibility score threshold of 0.1, and displayed scaled gene expression values from Seurat for the indicated differentially expressed genes linked to chamber-dependent distal DA cCREs.

### GREAT ontology analysis

The Genomic Regions Enrichment of Annotations Tool (GREAT, http://great.stanford.edu/public/html/index.php)^28^ was used with default settings for indicated cCREs or candidate enhancers in the .bed format. Biological process enrichments are reported. P-values shown for enrichment are Bonferroni-corrected binomial p-values.

### Motif enrichment analysis

For *de novo* and known motif enrichment analysis of cluster-specific cCREs, the *findMotifsGenome.pl* utility of the HOMER package was used with default settings^30^. For display of enrichment patterns for motifs from the JASPAR^103^ database with evidence of enrichment in at least one set of cell type-specific cCREs, motifs with an enrichment p- value < 10^-5^ in at least one set of cluster-specific cCREs were selected. For motif enrichment within differentially accessible cCREs, *narrowPeak* calls from MACS2 were used as input, with peaks called on the corresponding cell type (as described above) used as background. For enrichment of motifs within deconvoluted bulk enhancers, snATAC-seq peaks from the union of snATAC-seq peaks were utilized. Summits were extracted from peaks that overlapped bulk enhancer annotations and extended by 250bp on either side to obtain fixed-width peaks. We also computed motif enrichment scores at single-cell resolution using chromVAR^29^. For input to chromVAR, we used the summits of the 287,415 peaks in our consensus list extended by 250 base pairs in either direction, and a set of 870 non-redundant motifs as input. To identify differentially enriched motifs in each cell type, we used the following strategy: for each cell type and each motif, we computed a Rank Sum test between the chromVAR Z-score distributions from cells within the cell type and outside of the cell type. Tests were run using a random sampling of 40,000 cells. Then, for each cell type we used 1e-8 as p-value cutoff. In addition, we applied a Bonferroni correction to account for multiple testsing which resulted in selection of significant motifs with p-value < 1e-11.

### Bulk candidate heart enhancer deconvolution

We obtained published candidate heart enhancers annotated by H3K27ac ChIP-seq from a recently reported bulk survey of healthy left ventricular tissue from 18 human donors^14^. Candidate enhancers were defined per the study as H3K27ac ChIP-seq peaks that were at least 1kb away from a transcription start site and present in two or more donors. Because these reference annotations were derived from bulk profiling of healthy left ventricles, we selected only left ventricular nuclei from our aggregate dataset for comparison. We limited our analysis to cell types that comprised at least 5% of nuclei by proportion in our aggregate dataset. These included cardiomyocytes, fibroblasts, endothelial cells, smooth muscle cells, and macrophages. We first combined all fragments for each cell type from left ventricular datasets. The ‘coverage’ option of BEDtools^80^ was applied with default settings to count the total number of chromatin fragments from each ventricular cell type overlapping the candidate enhancer annotations. This yielded a raw count matrix in the format of snATAC-seq cell types (columns) by candidate enhancers (rows). The raw count matrix was normalized to RPKM (reads per kilobase per million mapped reads) for each candidate enhancer. We next used Cluster3.0^104^ to k-means cluster the 31,033 healthy heart candidate enhancers into K groups between 2 and 12 with the following settings (Method = *k-Means*, Similarity Metric = *Euclidian distance*, number of runs = *100*). We calculated the Davies-Bouldin (DB) index^81^ as described above for each clustering using the *index.DB* function of the R package clusterSim (http://keii.ue.wroc.pl/clusterSim/). We selected a k-means of 8, which yielded the lowest DB index, indicating the best partitioning.

We repeated the above analysis for 4,406 candidate enhancers reported have increased bulk H3K27ac ChIP signal and 3,101 candidate enhancers reported to have decreased signal in 18 late stage idiopathic dilated cardiomyopathy (heart failure) left ventricles versus 18 healthy control left ventricles reported in the same study. We again clustered the candidate enhancers for both groups into k groups between 2 and 12 as above and selected the clustering that yielded the lowest DB index^81^.

### Genome-wide association study (GWAS) variant enrichment analysis

We used LD (linkage disequilibrium) score regression^41, 105^ to estimate genome-wide enrichment for variants associated with GWAS traits within cell type-resolved open chromatin sites. We compiled published GWAS summary statistics for cardiovascular diseases^42–46^, other diseases^106–117^, and non-disease traits^118–127^ using the European subset from transethnic studies where applicable. We created custom LD score files by using peaks from each cluster as a binary annotation. In addition to the baseline annotations included in the baseline-LD model v2.2, we also included LD scores created from the merged peaks across all clusters as the background. For each trait, we used LD score regression to estimate enrichment z-scores for each annotation relative to the background. Using these z-scores, we computed one-sided p-values for enrichment and used the Benjamini Hochberg procedure to correct for multiple tests.

### Fine mapping for atrial fibrillation

We obtained published atrial fibrillation GWAS summary statistics and index variants for 111 disease-associated loci^43^. To construct credible sets of variants for each locus, we first extracted all variants in linkage disequilibrium (r^2^ > 0.1 using the EUR subset of 1000 Genomes Phase 3)^128^ in a large window (±2.5 Mb) around each index variant. We next calculated approximate Bayes factors^47^ (ABF) for each variant using effect size and standard error estimates. We then calculated posterior probabilities of association (PPA) for each variant by dividing its ABF by the sum of ABF for all variants within the locus. For each locus, we then defined 99% credible sets by sorting variants by descending PPA and retaining variants that added up to a cumulative PPA of > 0.99. This resulted in an output of 6,014 candidate causal variants.

### Variant prioritization for functional validation

To prioritize variants for functional validation, we refined our list of candidate causal variants from fine mapping analysis to only those with a posterior probability of association (PPA) > 0.1 (216 remaining out of 6,014). We used BEDtools^80^ to intersect these variants with ATAC-seq peaks called on an aggregate .bed file for atrial and ventricular cardiomyocyte snATAC-seq clusters (cardiomyocyte cCREs). This resulted in 40 fine-mapped variants that resided within 38 candidate cardiomyocyte cCREs (38 cCRE-variant pairs).

We assessed each remaining cCRE-variant pair via the following criteria:

- cCREs primarily accessible in cardiomyocytes
- presence of a corresponding ATAC-seq peak at a testable time point in the *in vitro* hPSC-cardiomyocyte differentiation model system
- sequence conservation in 100 vertebrates (genome browser track generated using phyloP of the PHAST5 package downloaded from UCSC genome browser^129^, http://hgdownload.soe.ucsc.edu/goldenPath/hg38/phyloP100way/)
- predicted co-accessibility of candidate enhancer with a gene promoter
- expression of putative target gene associated with cCRE appearance (chromatin accessibility and H3K27ac) during hPSC-cardiomyocyte differentiation^49^

A candidate cCRE-variant pair at the *KCNH2* locus was prioritized for functional experimentation.

### ChIP-seq data processing

Reads were mapped to the human genome reference GRCh38 using Bowtie2 (version 2.2.6)^130^ and reads with MAPQ > 30 selected using SAMtools (version 1.3.1)^131^. PCR duplicates were removed using MarkDuplicates function of Picard tools (version 1.119)^132^. RPKM normalized signal tracks were generated using BamCoverage function in deepTools (version 2.4.1)^133^.

### RNA-seq data processing

Reads were mapped to the human genome reference GRCh38 using STAR (version 020201)^134^ and reads with MAPQ > 30 selected using SAMtools (version 1.3.1)^131^. PCR duplicates were removed using MarkDuplicates function of Picard tools (version 1.1.19)^132^. RPKM normalized signal tracks were generated using BamCoverage function in deepTools (version 2.4.1)^133^.

### ATAC-seq data processing

Reads were mapped to the human genome reference GRCh38 using Bowtie2 (version 2.2.6)^130^ and reads with MAPQ > 30 selected using SAMtools (version 1.3.1)^131^. PCR duplicates were removed using SAMtools (version 1.3.1)^131^. RPKM normalized signal tracks were generated using BamCoverage function in deepTools (version 2.4.1)^133^.

### Statistics

No statistical methods were used to predetermine sample sizes. There was no randomization of the samples, and investigators were not blinded to the specimens being investigated. However, clustering of single nuclei based on chromatin accessibility was performed in an unbiased manner, and cell types were assigned after clustering. Low-quality nuclei and potential barcode collisions were excluded from downstream analysis as outlined above. Cluster-specificity at each cCRE was tested using edgeR^102^ as described above, with p-values corrected via the Benjamini Hochberg method. To identify differentially accessible sites between heart chambers and for each cell type, a likelihood ratio test was used, and the resulting p-value was corrected using the Benjamini Hochberg method. For significance of ontology enrichments using GREAT, Bonferroni-corrected binomial p values were used^28^. For significance testing of enrichment of *de novo* and known motifs, a hypergeometric test was used without correction for multiple testing^30^. For luciferase and qPCR data, we performed one-way ANOVA (ANalysis Of VAriance) analysis with post-hoc Tukey HSD (Honestly Significant Difference) using GraphPad Prism version 8.0.0 for Windows, GraphPad Software, San Diego, California USA, www.graphpad.com.

### External datasets

Cardiomyocyte differentiation: RNA-Seq, H3K27ac day 0 (hPSC); day 5 (cardiac mesoderm); and day 15 (primitive cardiomyocytes) were downloaded from GSE116862^49^. Signal tracks for heart H3K27ac ChIP-seq data were downloaded from https://portal.nersc.gov/dna/RD/heart/. List of candidate enhancers was downloaded from Supplemental tables^14^. H3K27ac ChIP-seq data for cardiomyocyte nuclei from non-failing donors (NF1) were downloaded from NCBI SRA BioProject ID PRJNA353755^135^.

### Code availability

The pipeline for processing snATAC-seq data is available as a part of the Taiji software: https://taiji-pipeline.github.io/

Custom code used for demultiplexing and downstream analysis for snATAC data is available here: https://gitlab.com/Grouumf/ATACdemultiplex/-/tree/master/ATACdemultiplex https://gitlab.com/Grouumf/ATACdemultiplex/-/blob/master/scripts/

The protocol for the custom set of motifs used with chromVAR^29^ can be found here: https://github.com/GreenleafLab/chromVARmotifs

### Data availability

Data will be deposited to dbGAP. Processed data can be explored using our publicly-available web portal including a UCSC cell browser (https://github.com/maximilianh/cellBrowser) and genome browser track viewer (IGV.js: https://github.com/igvteam/igv.js#igvjs): http://catlas.org/humanheart.

