## Supplemental_Table_I for "Cardiac Cell Type-Specific Gene Regulatory Programs and Disease Risk Association"

| Donor # | Age | Race | Gender | Height (cm) | Weight (kg) |
|---|---|---|---|---|---|
| 1 | 55 | White | F | 165 | 100 |
| 2 | 53 | White | M | 185 | 70 |
| 3 | 48 | Hispanic | M | 183 | 97 |
| 4 | 67 | White | F | 160 | 108 |
